# New metabolic alterations and predictive marker pipecolic acid in sera for esophageal squamous cell carcinoma

**DOI:** 10.1101/2021.03.31.437882

**Authors:** Lei Liu, Jia Wu, Minxin Shi, Fengying Wang, Haimin Lu, Jibing Liu, Weiqin Chen, Guanzhen Yu, Dan Liu, Jing Yang, Qin Luo, Yan Ni, Xing Jin, Xiaoxia Jin, Wen-Lian Chen

**Affiliations:** Department of Thoracic Surgery, The Affiliated Tumor Hospital of Nantong University, Nantong 226300, Jiangsu Province, China; Cancer Institute, Longhua Hospital Shanghai University of Traditional Chinese Medicine, Shanghai 200032, China; Department of Epidemiology, Tumor Institute, The Affiliated Tumor Hospital of Nantong University, Nantong 226300, Jiangsu Province, China; Department of Clinical Laboratory, Longhua Hospital Shanghai University of Traditional Chinese Medicine, Shanghai 200032, China; The Children’s Hospital, Zhejiang University School of Medicine, National Clinical Research Center for Child Health, Hangzhou 310029, China; Department of Pathology, The Affiliated Tumor Hospital of Nantong University, Nantong 226300, Jiangsu Province, China

**Author notes:** Corresponding authors. (Xing Jin), (Xiaoxia Jin), (Wen-Lian Chen). Equal contribution (Lei Liu, Jia Wu, Minxin Shi).

**Keywords:** Esophageal squamous cell carcinoma, Serum metabolome, Esophagectomy, Predictive potential, Pipecolic acid

## Abstract

Esophageal squamous cell carcinoma (ESCC) is a major histological subtype of esophageal cancer with dismal prognosis. Although several serum metabolomic investigations have been reported, ESCC tumor-associated metabolic alterations along with predictive biomarkers in sera were not defined. Here we enrolled 34 treatment-naive ESCC patients and collected their pre-and post-esophagectomy sera together with sera from 34 healthy volunteers for metabolomic survey. Our comprehensive analysis discerned ESCC tumor-associated metabolic alterations as represented by a panel of 12 serum metabolites. Notably, postoperative abrosia and parenteral nutrition significantly perturbed the serum metabolome. Furthermore, we performed examination using sera from carcinogen-induced mice at dysplasia and ESCC stages, and identified three ESCC tumor-associated metabolites conserved between mice and humans. Notably, among these metabolites, pipecolic acid was progressively increased in mouse sera from dysplasia to cancerization, and it could accurately discriminate between mice at dysplasia stage and healthy control mice. Furthermore, this metabolite was essential for ECSS cells to oppose oxidative stress-induced DNA damage and cell proliferation arrest. Together, this study uncovered 12 ESCC tumor-associated serum metabolites with potential for monitoring therapeutic efficacy and disease relapse, presented evidence for refining parenteral nutrition composition, and highlighted serum pipecolic acid as an attractive biomarker for prediction of ESCC tumorigenesis.

## Introduction

Esophageal cancer (EC) is the ninth most common cancer and the sixth leading cause of cancer death globally, and its incidence is still increasing steadily [1, 2]. Esophageal squamous cell carcinoma (ESCC) and esophageal adenocarcinoma are two main histological types of EC. In the highest-risk region from northern Iran through Central Asia to north-central China, ESCC is the most prevalent type accounting for 90% of EC, and patients with ESCC in China account for more than half of the global ESCC cases [2–5]. To date, esophagectomy is the primary treatment for initially diagnosed patients [6]. Chemotherapy or chemoradiotherapy is required for patients with locally advanced or cN1-N3 EC [2]. Due to lack of specific signs or symptoms at early stage, the majority of diagnosed ESCC patients are at advanced stage [7, 8]. Therefore, the prognosis and 5-year survival (15-20%) of these patients are unsatisfactory [9–11]. Early diagnosis is key for improving the therapeutic outcomes of ESCC patients. Hence, it is urgent to identify new biomarkers with predictive potential for ESCC tumorigenesis.

Metabolic reprogramming is a core hallmark of cancer [12, 13]. Neoplastic cells require a large amount of energy and raw materials in the process of cancer occurrence, development and metastasis, thus disturbing the global metabolism including metabolic signatures of circulating blood of patients [12–17]. Abnormal metabolites in peripheral blood can be used for cancer diagnosis, treatment efficacy assessment and patient prognosis prediction [15, 16, 18]. Hence, systemic measurement of metabolite composition in sera would deepen our understanding of ESCC molecular features and yield to new diagnostic chances for ESCC patients. Several metabolomic studies using serum or plasma specimens of ESCC patients have been conducted, and some significant variations in lipid, glucose, and amino acid metabolism in ESCC patients as compared to healthy controls have been identified [19–24]. However, due to metabolic modifications induced by ESCC indirectly as confounding factors in these data, ESCC tumor-associated metabolic alterations cannot be discerned. A recent study collected pre- and post-operative serum samples of ESCC patients and performed metabolomic investigation in order to identify diagnostic biomarkers [25]. Nevertheless, these pre-and post-operative serum samples were derived from different ESCC patient cohorts and the measurement platform nuclear magnetic resonance spectroscopy had limited resolution [25]. Therefore, this study cannot exactly and comprehensively capture ESCC tumor-associated metabolic alterations.

In this study, we enrolled 34 treatment-naive ESCC patients and 34 healthy volunteers. Pre- and post-esophagectomy sera from ESCC patients along with sera from healthy volunteers were collected for metabolomic investigation using gas chromatography-time-of-flight mass spectrometry (GC-TOFMS), aiming at discovering ESCC tumor-associated metabolic alterations. Subsequently, we recruited control mice and carcinogen-induced mice at dysplasia and ESCC stages. Sera of these mice were examined to validate ESCC tumor-associated metabolic alterations and to identify metabolite biomarkers with predictive potential. In the end, we conducted functional assays to explore the role of identified metabolite biomarker in ESCC tumorigenesis.

## Results

### A distinct serum metabolic signature of preoperative patients with ESCC

The design of this study was showed in **Figure 1**A. We harvested serum samples from 34 ESCC patients (Pre-esophagectomy versus post-esophagectomy) and healthy controls (HCs) to identify ESCC tumor-associated serum metabolites. Meanwhile, we collected serum samples from control mice and 4-nitroquinoline 1-oxide (4-NQO)-treated mice at dysplasia and ESCC stages to discover ESCC tumor-associated metabolites conserved between mice and humans, and to find serum metabolite markers for ESCC prediction. Finally, we performed functional assays for the predictive metabolite marker. There were no significant differences in age, gender, serum alanine aminotransferase, serum aspartate aminotransferase and serum creatinine between HCs and preoperative ESCC patients, or between preoperative and postoperative ESCC patients (**Table 1**). Before metabolomic data analysis, we firstly assessed data quality and observed low relative standard deviation values of chemical standards in quality control (QC) samples, revealing high stability of the measurement system (**Table S1**). A total of 161 metabolites were identified after excluding internal standards and unknown peaks in this study. The serum metabolomic feature of preoperative ESCC patients was remarkably altered as relative to that of HCs, as demonstrated by the robust model of orthogonal partial least squares-discriminant analysis (OPLS-DA, R^2^Y = 0.88, Q^2^ = 0.81) (Figure 1B). Among those 161 metabolites, 58 of them were significantly changed in preoperative ESCC sera (Bonferroni-adjusted *P* < 0.05), including 14 amino acids, 7 carbohydrates, 18 lipids, 2 nucleosides, 10 organic acids and 7 unclassified metabolites (Figure 1C). Furthermore, among these 58 changed metabolites, 16 of them were upregulated, including 4 amino acids, 1 carbohydrate, 3 lipids, 1 nucleoside, 5 organic acids and 2 unclassified metabolites; while the remaining metabolites were downregulated, including 10 amino acids, 6 carbohydrates, 15 lipids, 1 nucleoside, 5 organic acids and 5 unclassified metabolites. In agreement with previous studies [20, 25], our data revealed that glutamine was upregulated in ESCC sera, while glutamic acid, alanine, glycine, serine, ornithine, ribose, glyceric acid, docosahexaenoic acid and linoleic acid were restrained in ESCC sera.

**Table 1.**
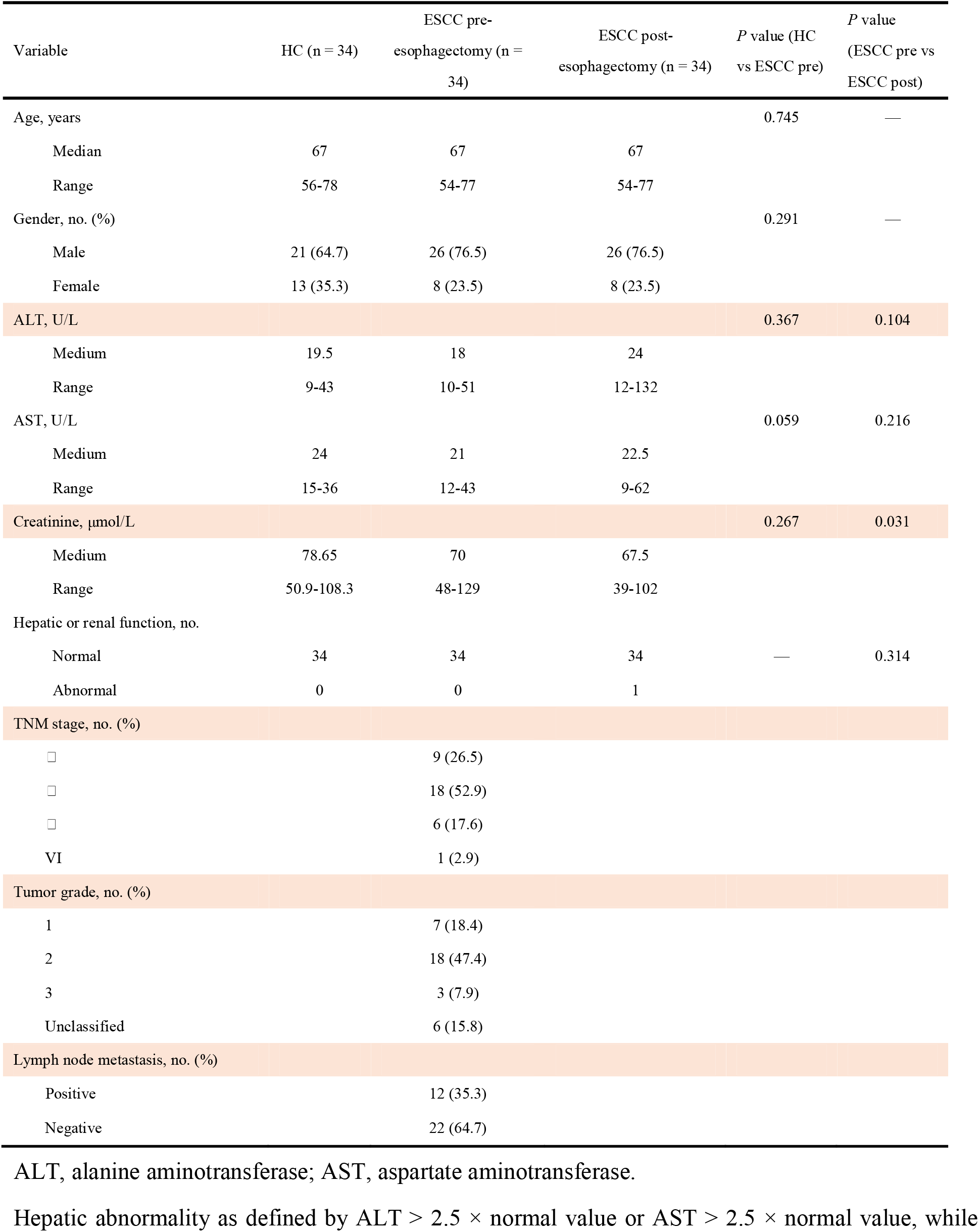
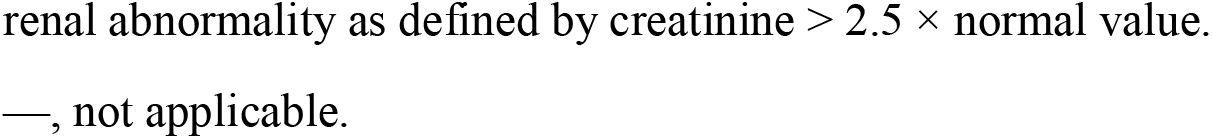
Basic characteristics of enrolled healthy volunteers and patients with ESCC.

| Variable | HC (n = 34) | ESCC pre-esophagectomy (n = 34) | ESCC post-esophagectomy (n = 34) | *P* value (HC vs ESCC pre) | *P* value (ESCC pre vs ESCC post) |
| --- | --- | --- | --- | --- | --- |
| Age, years |  |  |  | 0.745 | — |
| Median | 67 | 67 | 67 |  |  |
| Range | 56-78 | 54-77 | 54-77 |  |  |
| Gender, no. (%) |  |  |  | 0.291 | — |
| Male | 21 (64.7) | 26 (76.5) | 26 (76.5) |  |  |
| Female | 13 (35.3) | 8 (23.5) | 8 (23.5) |  |  |
| ALT, U/L |  |  |  | 0.367 | 0.104 |
| Medium | 19.5 | 18 | 24 |  |  |
| Range | 9-43 | 10-51 | 12-132 |  |  |
| AST, U/L |  |  |  | 0.059 | 0.216 |
| Medium | 24 | 21 | 22.5 |  |  |
| Range | 15-36 | 12-43 | 9-62 |  |  |
| Creatinine, $\mu\text{mol/L}$ | | | | 0.267 | 0.031 |
| Medium | 78.65 | 70 | 67.5 |  |  |
| Range | 50.9-108.3 | 48-129 | 39-102 |  |  |
| Hepatic or renal function, no. |  |  |  |  |  |
| Normal | 34 | 34 | 34 | — | 0.314 |
| Abnormal | 0 | 0 | 1 |  |  |
| TNM stage, no. (%) |  |  |  |  |  |
| I |  | 9 (26.5) |  |  |  |
| II |  | 18 (52.9) |  |  |  |
| III |  | 6 (17.6) |  |  |  |
| IV |  | 1 (2.9) |  |  |  |
| Tumor grade, no. (%) |  |  |  |  |  |
| 1 |  | 7 (18.4) |  |  |  |
| 2 |  | 18 (47.4) |  |  |  |
| 3 |  | 3 (7.9) |  |  |  |
| Unclassified |  | 6 (15.8) |  |  |  |
| Lymph node metastasis, no. (%) |  |  |  |  |  |
| Positive |  | 12 (35.3) |  |  |  |
| Negative |  | 22 (64.7) |  |  |  |

ALT, alanine aminotransferase; AST, aspartate aminotransferase.
Hepatic abnormality as defined by $\text{ALT} > 2.5 \times \text{normal value}$ or $\text{AST} > 2.5 \times \text{normal value}$ , while
renal abnormality as defined by creatinine $> 2.5 \times$ normal value.
—, not applicable.

**Figure 1.**
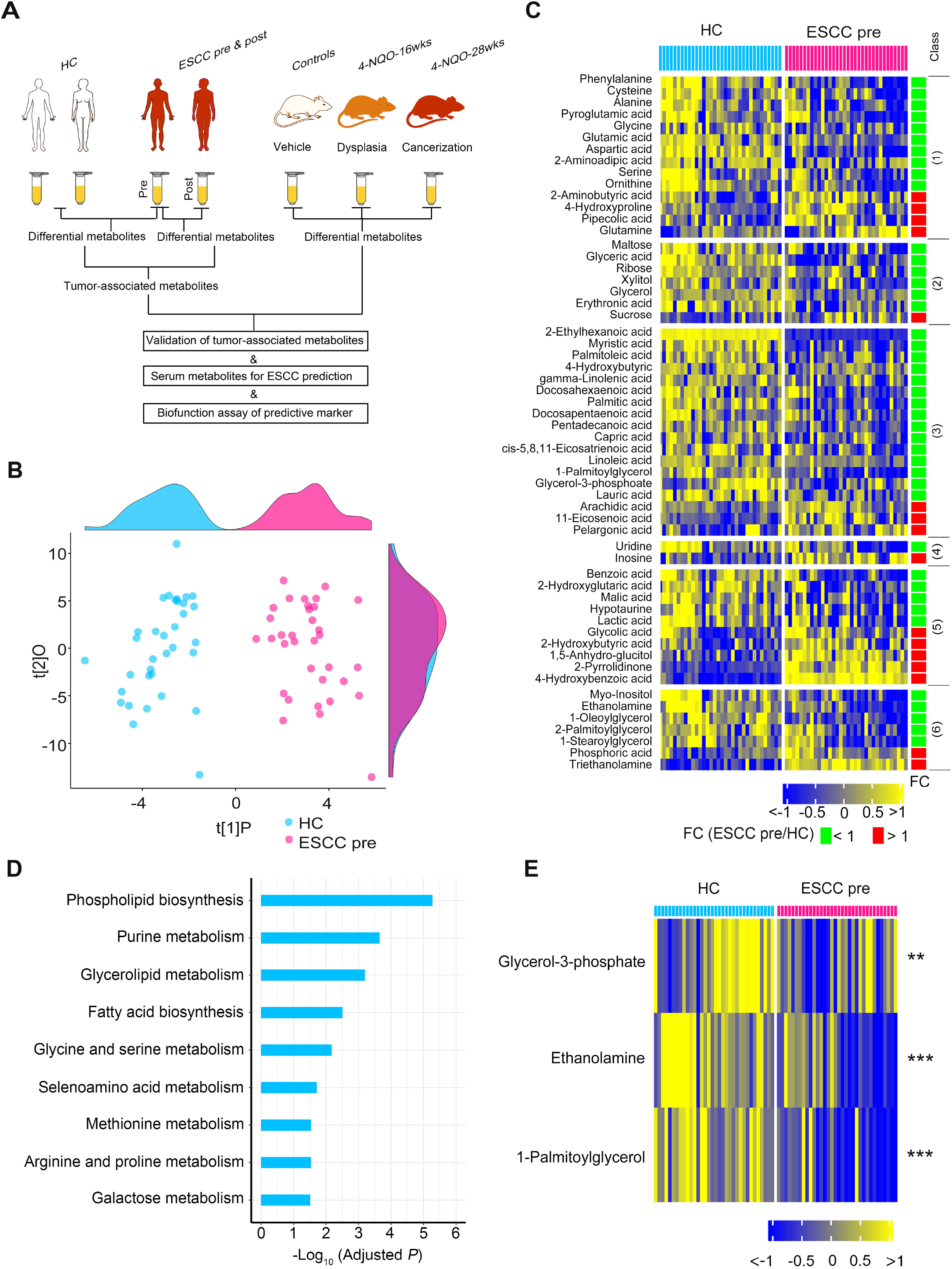
Altered serum metabolic signature of preoperative patients with ESCC. **A.** Study design. Serum samples from ESCC patients (Pre-esophagectomy versus post-esophagectomy) and healthy controls (HCs) were harvested for identification of ESCC tumor-associated serum metabolites. Meanwhile, serum samples from control mice and 4-nitroquinoline 1-oxide (4-NQO)-treated mice at dysplasia and ESCC stages were collected for discovery of ESCC tumor-associated metabolites conserved between mice and humans as well as serum metabolite markers for ESCC prediction. Finally, functional assays of the predictive metabolite marker would be conducted. Pre, Pre-esophagectomy; Post, Post-esophagectomy. **B.** OPLS-DA score plot showing a global metabolic difference in sera between preoperative ESCC patients (ESCC pre) and HCs. The density plots of principal component 1 and principal component 2 were displayed on the top and right-hand sides of the PCA score plots respectively. **C.** Heat map showing 58 differentially expressed serum metabolites in preoperative ESCC patients compared to HCs. The metabolites are subclassified as follows: (1) Amino acids, (2) Carbohydrates, (3) Lipids including fatty acids, (4) Nucleotides, (5) Organic acids, (6) Unclassified. **D.** Metabolic pathway enrichment analysis using differential metabolites between preoperative ESCC patients and HCs. **E.** Heat map showing the mostly disturbed metabolic pathway phospholipid biosynthesis in preoperative ESCC patients.

Next, we performed metabolite set enrichment analysis (MSEA) analysis using above 58 modified metabolites to observe which metabolic pathways were perturbed in preoperative ESCC sera. The result manifested that a total of 9 metabolic pathways were significantly altered (Holm-adjusted *P* < 0.05) (Figure 1D). For the mostly disturbed metabolic pathway phospholipid biosynthesis, all detected metabolites involved in this pathway, including glycerol-3-phosphate, ethanolamine and 1-palmitoylglycerol, were consistently declined in preoperative ESCC sera (Figure 1E).

### Identification of ESCC tumor-associated metabolic alterations in sera

As we stated in the Materials and Methods section, all enrolled ESCC patients received esophagectomy to completely remove tumors. Subsequently, these patients were given abrosia and persistent parenteral nutrition from day 1 after the surgery. At day 3 after esophagectomy, serum samples from these patients were harvested for metabolomic measurement. Firstly, we performed comparative analysis between preoperative and postoperative ESCC serum samples. By comparison with the preoperative sera, postoperative sera exhibited a remarkably modified metabolic profile as illustrated by the robust OPLS-DA model (R^2^Y = 0.84, Q^2^ = 0.76) (**Figure 2**A), demonstrating that esophagectomy together with postoperative abrosia and parenteral nutrition replenishment could result in an overt impact on the global metabolism of ESCC patients. A total of 78 metabolites were found to be significantly changed in postoperative ESCC sera as relative to preoperative specimens (Bonferroni-adjusted *P* < 0.05), including 16 amino acids, 16 carbohydrates, 18 lipids, 2 nucleosides, 19 organic acids and 7 unclassified metabolites (Figure 2B). Subsequently, we conducted MSEA analysis using these 78 metabolites and found that 10 metabolic pathways were significantly fluctuated in postoperative ESCC sera when compared to preoperative ESCC sera (Holm-adjusted *P* < 0.05) (Figure 2C). Notably, the mostly disturbed pathway was galactose metabolism. At metabolite level, 4 of 5 metabolites involved in this pathway were raised, while the remaining one was restrained in postoperative sera (Figure 2D).

**Figure 2.**
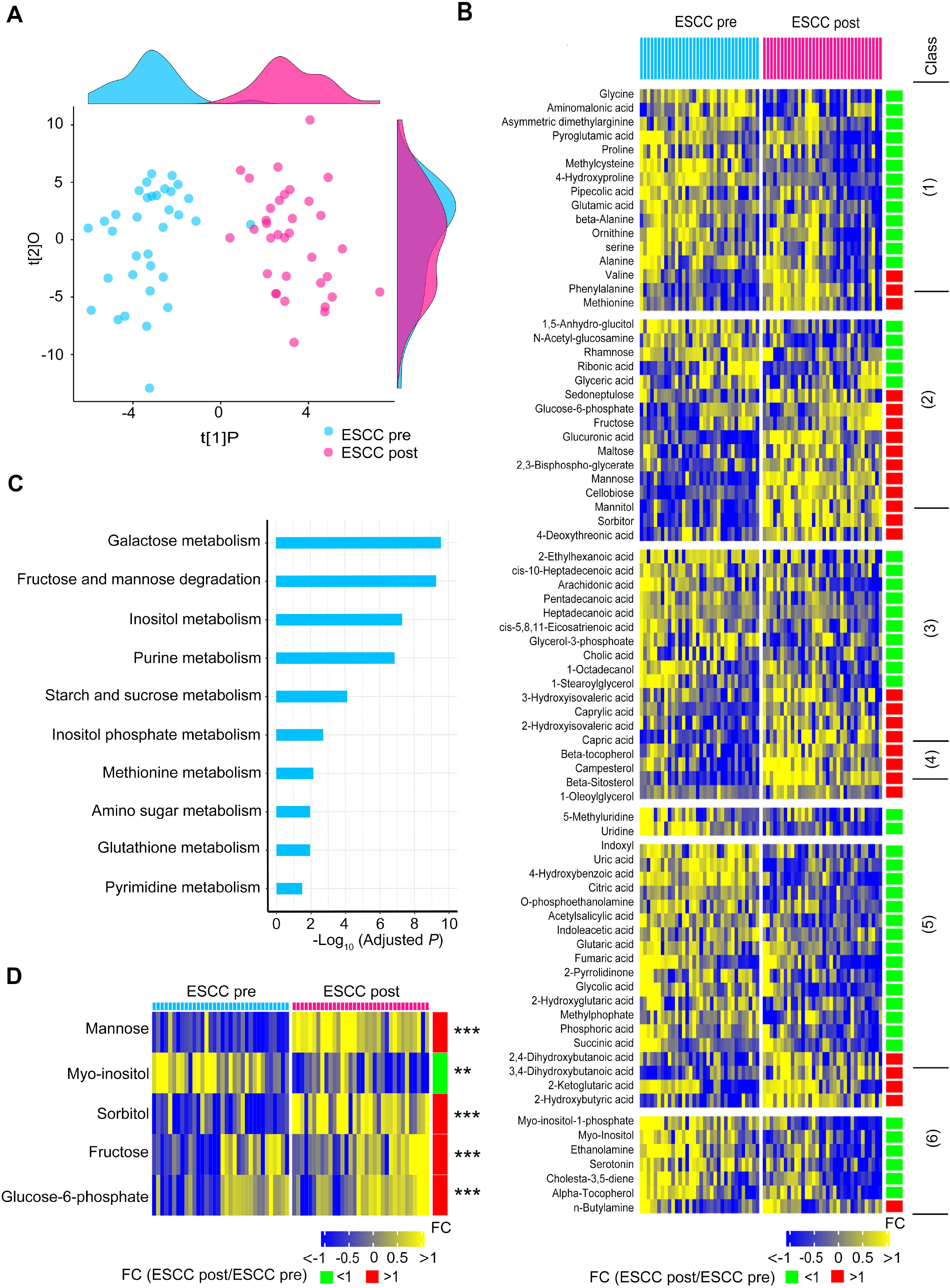
Serum metabolic alterations between preoperative and postoperative patients with ESCC. **A.** OPLS-DA score plot showing modified global metabolism in postoperative ESCC patients (ESCC post) compared to preoperative patients (ESCC pre). The density plots of principal component 1 and principal component 2 were displayed on the top and right-hand sides of the OPLS-DA score plots respectively. **B.** Heat map exhibiting 78 differential metabolites in postoperative ESCC patients relative to preoperative ESCC patients. The metabolites are subclassified as follows: (1) Amino acids, (2) Carbohydrates, (3) Lipids including fatty acids, (4) Nucleotides, (5) Organic acids, (6) Unclassified. **C.** Metabolic pathway enrichment analysis using differential metabolites between preoperative and postoperative ESCC patients. **D.** Heat map showing the mostly disturbed metabolic pathway galactose metabolism in postoperative ESCC patients. ^**^*P* < 0.01, ^****^*P* < 0.0001.

To determine the ESCC tumor-associated metabolic alterations, we performed a comprehensive analysis using data of HC, preoperative and postoperative ESCC groups. Logically, 58 differential metabolites in preoperative ESCC sera as compared to HC sera (figure 1B) contained metabolic signals altered by ESCC directly and indirectly. Therefore, we selected these metabolites to extract ESCC tumor-associated metabolic alterations. As shown in **Figure 3A**, 12 metabolites in postoperative ESCC sera were significantly restored toward HC direction. This indicated that these serum metabolites were specifically modulated by ESCC tumors of patients. These metabolites were enriched in 2 pathways, including ubiquinone biosynthesis and phenylalanine and tyrosine metabolism (Holm-adjusted *P* < 0.05) (Figure 3B). In addition, the heat map of Figure 3C exhibited that the remaining 46 metabolites in postoperative ESCC sera were changed toward an opposite HC direction or were not significantly altered as relative to preoperative ESCC sera. We concluded that these 46 metabolites were not specifically modulated by ESCC tumors. Of note, these metabolites were enriched in 8 pathways (Holm-adjusted *P* < 0.05) (Figure 3D).

**Figure 3.**
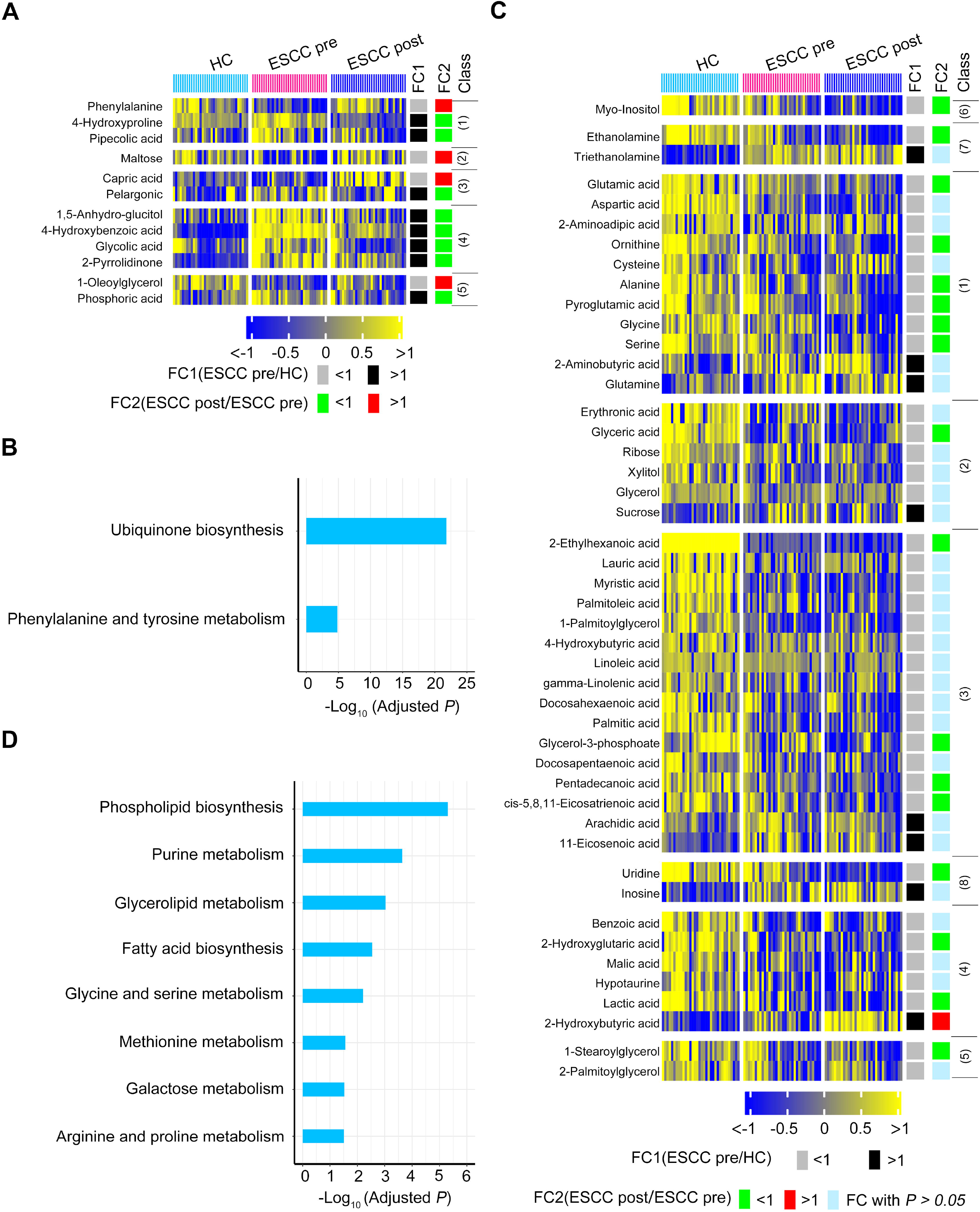
Alteration of 58 differential serum metabolites between preoperative patients and HCs in postoperative patients. **A, B.** Twelve metabolites in postoperative ESCC sera significantly restored toward HC direction (A), and metabolic pathway enrichment analysis of these metabolites (B). **C, D.** The remaining 46 metabolites in postoperative ESCC sera changed toward an opposite HC direction or not significantly altered as relative to preoperative ESCC sera (C), and metabolic pathway enrichment analysis of these metabolites (D). These metabolites were classified as follows (1) Amino acids, (2) Carbohydrates, (3) Lipids including fatty acids, (4) Organic acids, (5) Others, (6) Alcohols, (7) Amines, (8) Nucleotides.

### Metabolic perturbations caused by postoperative abrosia and parenteral nutrition

There were 58 differential metabolites in preoperative ESCC sera as relative to HC sera, whereas there were 78 differential metabolites in postoperative ESCC sera as relative to preoperative ESCC sera. This suggested that postoperative abrosia and parenteral nutrition might overtly disturb the serum metabolic profile of ESCC patients. To verify this speculation, we analyzed those 108 serum metabolites that were not altered in preoperative ESCC as relative to HC. As shown in **Figure 4**A, 27 of these metabolites (25.00%) were significantly declined in postoperative ESCC as relative to preoperative ESCC or HCs. These decreased metabolites included 4 amino acids, 3 carbohydrates, 5 lipids, 1 nucleotide, 10 organic acids and 4 unclassified metabolites. Furthermore, 22 of 108 metabolites (20.37%) were remarkably increased in postoperative ESCC as relative to preoperative ESCC or HC (Figure 4B). These upregulated metabolites included 2 amino acids, 10 carbohydrates, 6 lipids, 3 organic acids and 1 unclassified metabolite.

**Figure 4.**
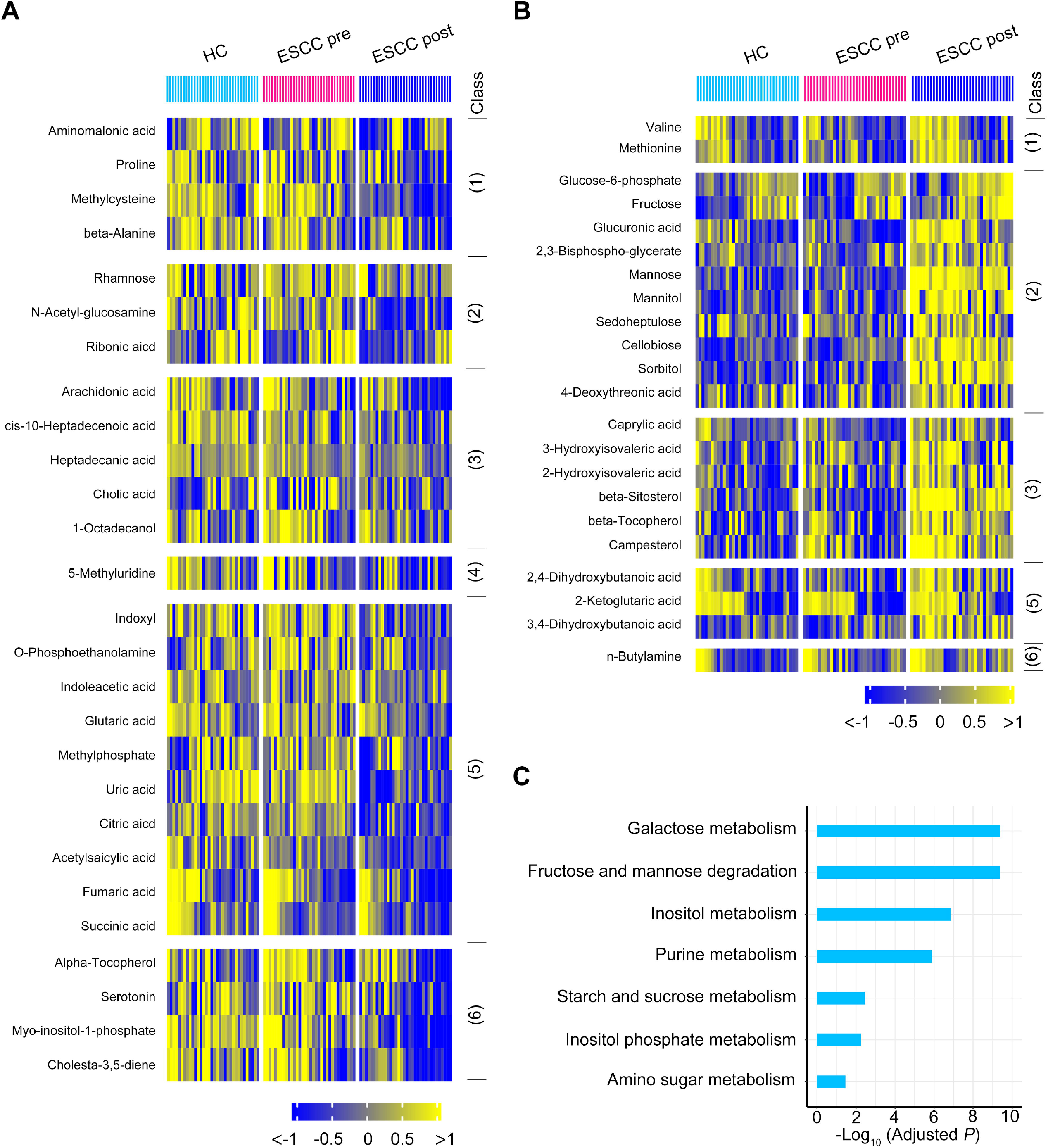
Metabolic fluctuation induced by postoperative abrosia and parenteral nutrition. **A, B.** For the 108 serum metabolites not altered between preoperative ESCC (ESCC pre) and HCs, 27 of them (25.00%) were significantly downregulated in postoperative ESCC (ESCC post) as relative to ESCC pre or HCs (A), while 22 of them (20.37%) were remarkably increased in ESCC post as relative to ESCC pre or HCs (B). These 49 metabolites are classified as follows: (1) Amino acids, (2) Carbohydrates, (3) Lipids including fatty acids, (4) Nucleotides, (5) Organic acids, (6) Unclassified. **C.** Metabolic pathway enrichment analysis of above 49 metabolites.

MSEA analysis using above 49 altered metabolites in sera of postoperative ESCC displayed that these compounds were enriched in 7 metabolic pathways (Figure 4C). Notably, carbohydrate anabolism was dramatically stimulated as shown by the upregulation of a total of 10 carbohydrates (Figures 4B and 4C). Collectively, among the 108 serum metabolites that were not modified between preoperative ESCC and HC, 45.37% of them were significantly perturbed in postoperative ESCC sera, indicating that postoperative abrosia and parenteral nutrition were able to result in an overt fluctuation of serum metabolome.

### Progressively modified serum metabolic signature of carcinogen-induced mice from dysplasia to ESCC

To validate the serum metabolic signature of preoperative ESCC patients and to discover metabolite biomarkers for prediction of ESCC tumorigenesis, we employed a carcinogen-induced ESCC mouse model as reported previously [26]. Mice were administered drinking water containing 100 μg/mL carcinogen 4-nitroquinoline 1-oxide (4-NQO) or propylene glycol (PG) as vehicle for 16 weeks, and then were followed-up for 12 weeks with pure drinking water. Thus, this gave rise to 3 groups of mice in this study, including control, 4-NQO-16wks and 4-NQO-28wks groups (**Figure 5**A). The disease progression was monitored by hematoxylin and eosin (H&E) staining of esophagi and by immunohistochemistry (IHC) assays for ESCC molecular biomarker keratin 14 (K14) along with cell proliferation marker Ki67 (Figure 5B). At week 16, esophageal squamous epithelia of 4-NQO group showed increased squamous basal cell layers, thickened spinous cells and disordered cell arrangement in each layer (Figure 5B, H&E image). Additionally, esophagi of 4-NQO-16-wks group revealed a feature of cell atypia characterized by an increase in cell size and ratio of nucleo-plasma, and by the nuclear hyperchromatism and pathological nuclear fission (Figure 5B, H&E image). Furthermore, the expression of K14 and Ki67 was enhanced and extended to the superficial cells as well as heteromorphic large cells (Figure 5B, IHC images). Together, these abnormal features demonstrated that mice of 4-NQO-16wks group were at dysplasia stage. At week 28, esophageal squamous epithelia of 4-NQO group exhibited high grade dysplasia, cancerization and microstromal infiltration (Figure 5B, H&E image). Notably, cells in the entire esophageal epithelial layer showed severe atypia. Additionally, heterotypical cells among these cells locally invaded into basement membranes with invasive growth (Figure 5B, H&E image). Moreover, the expression of K14 and Ki67 was obviously diffuse and highly strong (Figure 5B, IHC images). Collectively, these neoplastic characteristics indicated that mice of 4-NQO-28wks group were at cancerization stage.

**Figure 5.**
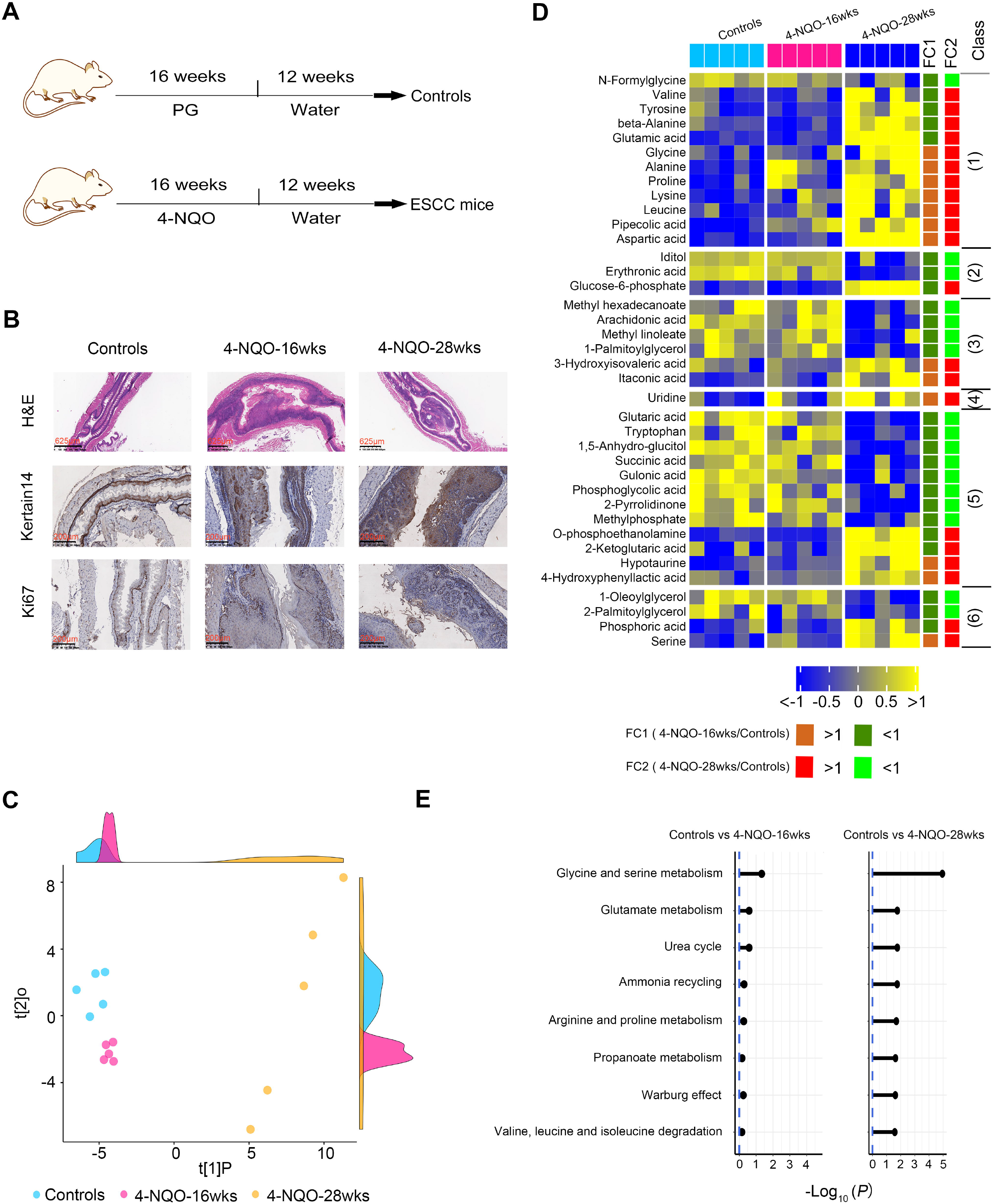
Progressively modified serum metabolome of carcinogen-induced mice from dysplasia to ESCC. A. scheme depicting 4-NQO induced ESCC mouse model from dysplasia to ESCC. PG, propylene glycol; 4-NQO, 4-nitroquinoline 1-oxide. B. Representative images of H&E staining and IHC staining of esophageal tissues of control mice and 4-NQO-treated mice. C. PLS score plot of serum metabolic profiles of control mice, 4-NQO-induced mice at dysplasia stage (4NQO-16wks) and 4-NQO-induced mice at cancerization stage (4NQO-28wks). The density plots of principal component 1 and principal component 2 were displayed on the top and right-hand sides of the PLS score plot respectively. **D, E.** Heat map displaying progressive changes of serum metabolites of mice from healthy control to cancerization stage (D) and metabolic pathway enrichment analysis of these metabolites (E).

When compared to the control group, both 4-NQO-16wks and 4-NQO-28wks groups showed obvious alteration for their serum metabolic profiles as illustrated by the robust model of partial least-squares regression (R^2^Y = 0.98, Q^2^ = 0.80) (Figure 5C). A total of 61 serum metabolites were significantly disturbed among these three groups. Notably, 38 of these disturbed metabolites were progressively modified from dysplasia stage to cancerization stage as compared to control group (Figure 5D), while the remaining 23 disturbed metabolites did not show this trend from normal control to cancerization (**Figure S1**).

Subsequently, we used above 38 progressively modified metabolites in Figure 5D to conduct MSEA analysis. The result manifested that 11 pathways were significantly disturbed in mouse sera at cancerization stage as compared to controls (Holm-adjusted *P* < 0.05) (Figure 5E). Notably, these metabolic pathways were slightly altered in mouse sera at dysplasia stage (Figure 5E).

### ESCC tumor-associated serum metabolites conserved between mice and humans with predictive potential for tumorigenesis

Subsequently, we wondered whether there were ESCC tumor-associated serum metabolites conserved between mice and humans with predictive potential for tumorigenesis. By comprehensively analyzing serum metabolomic data from ESCC mice and patients, we identified three metabolites conserved between mice and humans. Firstly, pipecolic acid was progressively increased in mouse sera from normal control, to dysplasia and to cancerization (**Figure 6A**). Consistently, this serum metabolite was dramatically elevated in preoperative patients but downregulated in postoperative cases (Figure 6B). Next, 1-oleoylglycerol was progressively reduced in mouse sera from normal control to cancerization (Figure 6C). In line with this finding, this serum metabolite was overtly suppressed in preoperative patients but upregulated in postoperative cases (Figure 6D). In addition, phosphoric acid in mouse sera was not disturbed at dysplasia stage as compared to control, whereas remarkably elevated at cancerization stage (Figure 6E). Correspondingly, this metabolite was markedly raised in preoperative patients but declined in postoperative cases (Figure 6F).

**Figure 6.**
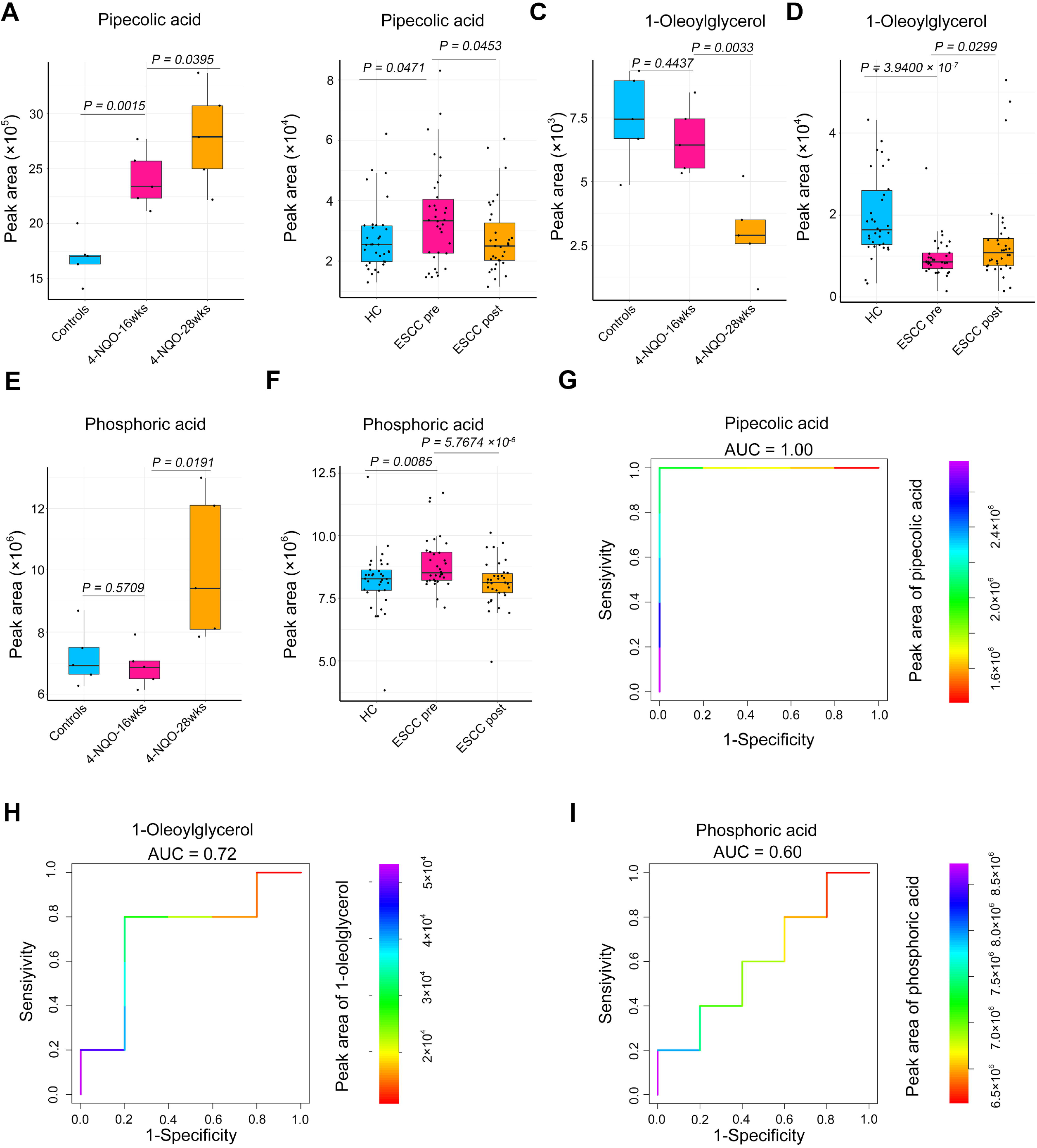
ESCC tumor-associated serum metabolites conserved between mice and humans with predictive potential for carcinogenesis. A. Abundance of serum pipecolic acid among control mice (n = 5), 4-NQO-induced mice at dysplasia stage (4NQO-16wks) (n = 5) and 4-NQO-induced mice at cancerization stage (4NQO-28wks) (n = 5). B. Abundance of serum pipecolic acid among HCs (n =34), preoperative ESCC patients (ESCC pre) (n = 34) and postoperative ESCC patients (ESCC post) (n = 34). C. Abundance of serum 1-oleoylglycerol among control mice (n = 5), 4-NQO-induced mice at dysplasia stage (4NQO-16wks) (n = 5) and 4-NQO-induced mice at cancerization stage (4NQO-28wks) (n = 5). D. Abundance of serum 1-oleoylglycerol among HCs (n =34), preoperative ESCC patients (ESCC pre) (n =34) and postoperative ESCC patients (ESCC post) (n =34). E. Abundance of serum phosphoric acid among control mice (n = 5), 4-NQO-induced mice at dysplasia stage (4NQO-16wks) (n = 5) and 4-NQO-induced mice at cancerization stage (4NQO-28wks) (n = 5). F. Abundance of serum phosphoric acid among HCs (n =34), preoperative ESCC patients (ESCC pre) (n = 34) and postoperative ESCC patients (ESCC post) (n = 34). **G-I.** ROC curves exhibiting the diagnostic capabilities of serum pipecolic acid (G), 1-oleoylglycerol (H) and phosphoric acid (I) to discriminate between healthy control (n =5) and precancerous mice (n = 5). Colors indicated metabolite abundance, and blue and red colors represented high and low abundance of metabolites respectively.

To ascertain whether above serum metabolites possessed predictive potential to accurately discriminate between healthy and precancerous cases, we performed receiver operating characteristic (ROC) curve analysis using control mice and 4-NQO-induced mice with dysplasia of esophageal epithelia. The values of area under the curves (AUC) were 1.00, 0.72 and 0.60 for pipecolic acid, 1-oleoylglycerol and phosphoric acid respectively, demonstrating that pipecolic acid possessed an excellent potential for predicting ESCC carcinogenesis (Figure 6G-6I).

### The protective effect of the predictive biomarker pipecolic acid on ESCC cells against oxidative stress

Finally, we explored whether the new biomarker identified in this study, pipecolic acid, possessed tumor-promoting role. It is known that elevated oxidative stress is extensively found in cancers, and the sustained rise of reactive oxygen species (ROS) promotes cell growth and facilitates carcinogenesis [27, 28]. However, overaccumulation of oxidative stress, once exceeding to the toxic threshold, would cause damage to lipids, proteins and DNA, and induce cell death [28, 29]. Hence, cancer cells require efficient antioxidation systems to limit oxidative stress overaccumulation, thus supporting cell survival and growth [28]. It is reported that pipecolic acid is able to enhance the anti-oxidant capability of mammalian cells [30]. Therefore, we assumed that increased pipecolic acid in the sera of preoperative patients and 4-NQO-treated mice at dysplasia and cancerization stages could enhanced the capability of ESCC cells to oppose oxidative stress in the tumor microenvironment. Rosup is a well-known compound mixture used for ROS generation [31]. Hence, Rosup was used to induce ROS in two human ESCC cells Eca109 and KYSE150 in the current study. Additionally, H_2_O_2_, one of ROS, was used as positive control.

We observed that addition of pipecolic acid dramatically palliated ROS production induced by Rosup and H_2_O_2_ in ESCC cells (**Figure 7A and B**). Moreover, addition of pipecolic acid overtly reversed DNA damage induced by Rosup and H_2_O_2_ as shown by the decreased expression of an established DNA damage marker phosphorylated H2AX (γ-H2AX) and by the downregulation of an DNA oxidative damage marker 8-oxo-7,8-dihydro-20-deoxyguanosine (8-oxo-dG) (Figure 7C-F). Consequently, pipecolic acid administration restored ESCC cell growth impaired by Rosup and H_2_O_2_ treatment in a dose-dependent manner (Figure 7G and H). In conclusion, pipecolic acid was essential for ESCC cells to sustain redox homeostasis and to antagonize oxidative stress-induced DNA damage and cell proliferation arrest.

**Figure 7.**
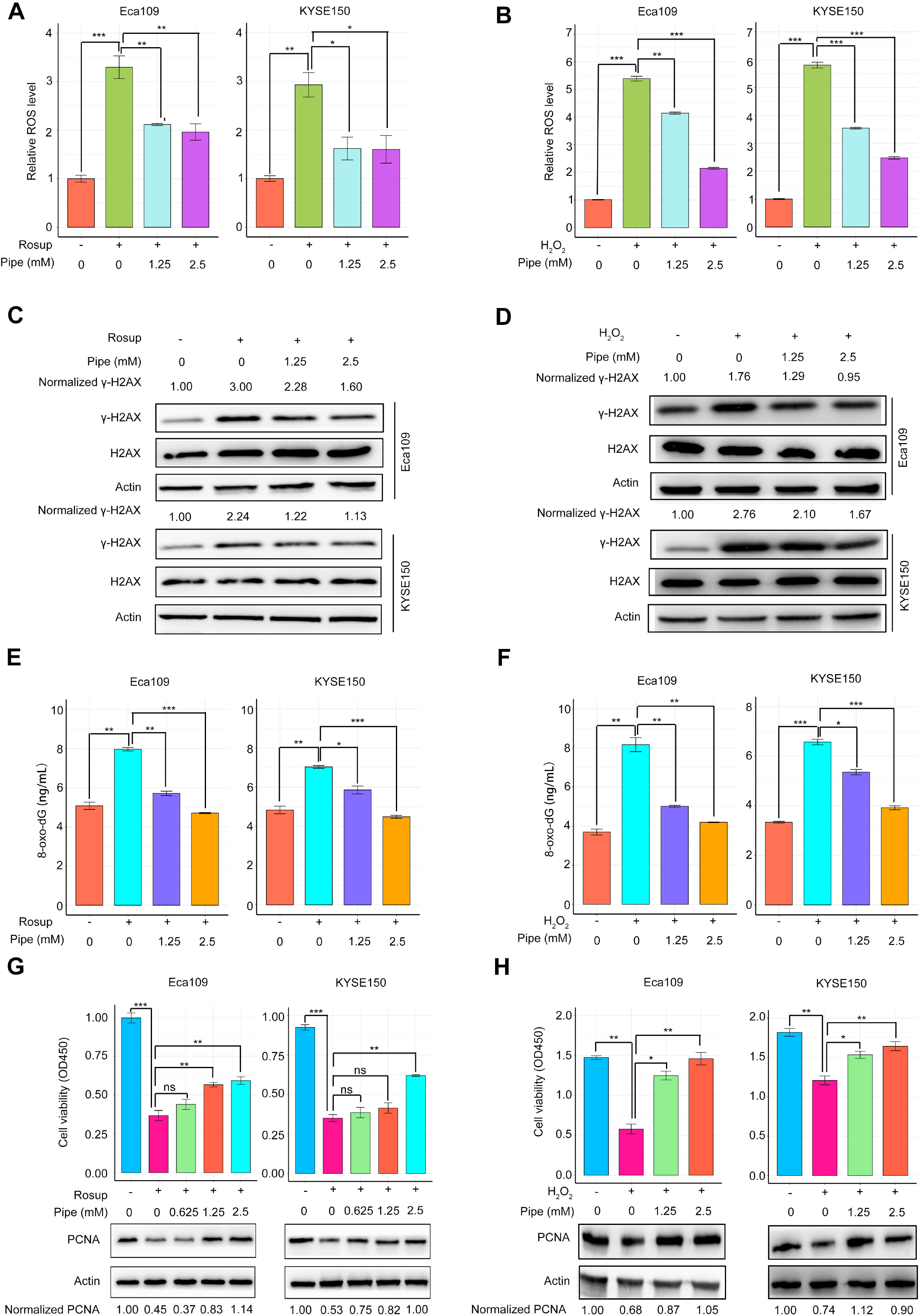
The antioxidant activity of the predictive biomarker pipecolic acid for ESCC cells. **A, B.** The impact of pipecolic acid (Pipe) on ROS production induced by Rosup (A) or H_2_O_2_ (B) in Eca109 and KYSE150 cells. **C, D.** The influence of Pipe on the expression of a DNA damage marker γ-H2AX induced by Rosup (C) or H_2_O_2_ (D) in Eca109 and KYSE150 cells. **E, F.** The influence of Pipe on the production of a DNA oxidative damage marker 8-oxo-dG induced by Rosup (E) or H_2_O_2_ (F) in Eca109 and KYSE150 cells. **G, H.** The impact of Pipe on the proliferation and PCNA expression of Eca109 and KYSE150 cells treated with Rosup (G) or H_2_O_2_ (H). Error bars represent mean ± SEM. ^*^*P* < 0.05, ^**^*P* < 0.01, ^***^*P* < 0.001, 2-tailed Student’s t test.

Circulating system is a crucial pathway for distal metastasis of cancer cells [32]. Due to pipecolic acid as a serum-containing metabolite in our study, we wondered whether it could impact the metastatic ability of ESCC cells. Treatment with pipecolic acid did not alter the migratory tendency as well as the expression of protein markers of epithelial-mesenchymal transition (EMT, an essential mechanism to drive cell migration) of ESCC cells (**Figure S2**), demonstrating that this metabolite did not perturb the metastatic ability of these malignant cells.

For another potential predictive marker 1-oleoylglycerol identified in Figure 6, we speculated that it had potential anti-cancer activity since that this metabolite was negatively associated with the existence of ESCC tumors. Indeed, administration of 100 μM 1-oleoylglycerol significantly restrained ESCC cell growth (**Figure S3**A and B). However, administration of 100 μM this metabolite did not elevate intracellular ROS generation, indicating that it repressed ESCC cell growth not via raising oxidative stress (Figure S3C and D).

## Discussion

As we stated in the introduction section, previous reports of ESCC serum metabolome are case-control studies [19–24]. A recent serum metabolomic study used pre- and post-operative serum samples of ESCC patients, whereas the pre- and post-operative patients belonged to distinct cohorts [25]. Thereby, ESCC tumor-associated metabolic alterations in sera could not be exactly extracted from these published data. In the current study, we collected pre- and post-esophagectomy sera from a same ESCC patient cohort along with sera from healthy volunteers to facilitate the identification of ESCC tumor-associated metabolic changes. A total of 58 metabolites were found to be dramatically disturbed in sera of preoperative ESCC patients compared to HCs. Importantly, our comprehensive analysis revealed that 12 of these serum metabolites were specifically linked to ESCC tumors, hence providing new biomarkers with potential for monitoring therapeutic efficacy and disease relapse. Of note, these 12 metabolites were enriched in ubiquinone biosynthesis and phenylalanine and tyrosine metabolism. A recent study finds that ubiquinone is essential for the growth of colon cancer organoids with p53 deficiency [33]. In our study, the biosynthetic precursor of ubiquinone biosynthesis, 4-hydroxybenzoic acid, was remarkably elevated in preoperative ESCC sera with a fold change of 4.30 (Figure 3A), indicating that this pathway was enhanced in ESCC patients. Dietary phenylalanine and tyrosine are required for the metastasis of melanoma, lung cancer and hepatocarcinoma cells [34]. The substrate of phenylalanine and tyrosine metabolism, phenylalanine, was restrained in preoperative ESCC sera with a fold change of −1.28 (Figure 3A), suggesting that this pathway was expedited in ESCC patients. However, the role of ubiquinone biosynthesis and phenylalanine and tyrosine metabolism in ESCC awaits to be elucidated.

An intriguing finding of the current study is the metabolic perturbations in sera of postoperative ESCC patients caused by administration of abrosia and parenteral nutrition. A panel of carbohydrates were markedly upregulated in postoperative sera, while there was only one known carbohydrate, glucose, in the parenteral nutrition solution. Of note, there were only two amino acid increased in postoperative sera, while there were 20 amino acids in the parenteral nutrition solution (**Table S2**). Together, these results indicated that postoperative ESCC patients required substantial carbohydrates and the components in parenteral nutrition solution were possibly used for conversion to these metabolites. Two essential amino acids, valine and methionine, are crucial for pancreatic cancer growth and lung cancer cell stemness respectively [35, 36]. These two amino acids were in parenteral nutrition solution and remarkably elevated in postoperative ESCC sera. Thus, it would be reasonable to reduce the amount of these two amino acids in parenteral nutrition solution in order to avoid activation of minimal residual disease. Moreover, it would be interesting to comprehensively ascertain the impact of metabolic perturbations induced by administration of postoperative abrosia and parenteral nutrition on the therapeutic outcomes of ESCC patients.

By enrolling a carcinogen-induced ESCC mouse model into the present study, we were able to delineate progressively modified serum metabolic signature from normal stage to cancerization stage. Furthermore, ESCC tumor-associated serum metabolites conserved between mice and humans were discovered, including pipecolic acid, 1-oleoylglycerol and phosphoric acid. To the best of our knowledge, there are no predictive biomarkers for ESCC carcinogenesis. Our result demonstrated that serum pipecolic acid is a potential biomarker with predictive potential for ESCC tumorigenesis as characterized by its high accuracy to discriminate between mice at dysplasia stage and control mice. Pipecolic acid has been reported to be able to elevate the anti-oxidant capability of mammalian cells [30]. In line with this previous finding, we found that this metabolite was able to oppose oxidative stress-induced DNA damage and cell growth arrest, indicating that this metabolite was potentially involved in ESCC tumorigenesis. Another interesting metabolite is 1-oleoylglycerol, which is negatively linked to ESCC tumorigenesis. We found that this metabolite impeded ESCC cell growth not via upregulating intracellular ROS.

Admittedly, there are several limitations in the current GC-TOFMS based study. Firstly, the ESCC tumor-associated metabolic alterations in sera should be validated in independent patient cohorts from other cancer centers. Secondly, the predictive potential of serum pipecolic acid for ESCC tumorigenesis should be verified using sera from patients with squamous dysplasia of esophageal epithelia. Additionally, metabolomic investigation of spent media of *in vitro* ESCC cell lines and *ex vivo* fresh ESCC tissues from patients should be performed to determine whether those abnormal serum metabolites discovered in this study are derived from ESCC cells.

## Materials and methods

### Enrollment of patients and healthy volunteers and collection of serum samples

A total of 34 patients with no prior treatment for their disease were enrolled into this study from the Affiliated Tumor Hospital of Nantong University (Nantong, Jiangsu Province, China). All patients were diagnosed as ESCC by histopathologic examination and staged according to the American Joint Committee on Cancer 8^th^ edition staging system [37]. From day 1 after esophagectomy, all cases were given abrosia and intravenous parenteral nutrition at a dosage of 3,000 mL/day. One dosage of parenteral nutrition contained 1,000 mL 5% glucose solution, 1,000 mL physiological saline, 500 mL 10% fat emulsion and 500 mL 10% amino acid solution (Table S2). In addition, age- and gender-matched healthy volunteers were enrolled from Longhua Hospital (Shanghai, China).

Overnight fasting peripheral blood samples were collected in the morning from healthy volunteers and ESCC patients before esophagectomy and after 3 days of surgery, and then transferred into vacuum blood collection tubes without any anticoagulants. All blood samples were clotted at room temperature for < 2 hours and centrifuged at 1,500 ×g for 10 minutes. Serum samples were obtained and stored at −80°C until analysis.

This study was approved by the Institutional Review Board (Approval number 2019-022) (**Figure S4**A). All participants provided written informed consent in accordance with the Declaration of Helsinki.

### Carcinogen-induced ESCC mice and sample harvest

We purchased six-week-old C57BL/6 female mice and used a reported chemical carcinogen 4-nitroquinoline 1-oxide (4-NQO, cat#N8141, Sigma, St Louis, MO) to induce ESCC as previously reported [38]. The study was approved and conducted by Wuhan Servicebio Technology Co., Ltd (Approved number 2018015) (Figure S4B). Briefly, 4-NQO stock solution was prepared in propylene glycol at a concentration of 6 mg/mL and then divided into aliquots for −20 □ storage. Two groups of mice were recruited. The experimental group was provided with the chemical carcinogen via drinking water containing 4-NQO with a final concentration of 100 μg/mL. While the control group was supplied with drinking water with no 4-NQO. Two groups received the same volume of propylene glycol in the drinking water, and drinking water was changed once a week. During the treatment, the mice were allowed to drink water whenever they wanted.

After a 16-week carcinogen or vehicle treatment, five control mice and five 4-NQO-induced mice were randomly selected and fasted overnight. In the next morning, these mice were killed via cervical dislocation, and their serum specimens and esophageal tissues were collected. Serum samples were quickly moved into −80 ◻ refrigerator for storage, while esophageal tissues were either flash frozen in liquid nitrogen or formalin-fixed paraffin-embedded and H&E stained.

Notably, the remaining five 4-NQO-induced mice were followed-up for another 12 weeks. At the end of the experiment, these mice were fasted overnight and killed in the next morning. Serum specimens and esophageal tissues were harvested and treated using a same protocol for the previous batch of mice. Moreover, immunohistochemistry staining assay were conducted for all paraffin-embedded tissues to measure the expression of ESCC biomarker keratin 14 and cell proliferation marker Ki67.

### Metabolomic profiling with GC-TOFMS

A total of 102 human sera together with 15 mouse sera were enrolled for metabolomic profiling using the GC-TOFMS platform. Metabolite extraction and measurement were executed using our previous protocol [15, 39–41]. Briefly, a 100 μL aliquot of each sample was used for metabolite extraction by adding 300 μL organic mixture (methanol:chloroform = 3:1, vol/vol). The mixture of organic solvent and serum was vortexed for 30 seconds, stored at −20 ◻ for 10 min, and centrifuged at 10,000 × g for 10 min. Subsequently, a 300 μL supernatant was transferred to a GC vial, spiked with internal standards including 10 μL of heptadecanoic acid (1 mg/mL) and 4-chlorophenylalanine (0.3 mg/mL), and then dried at −20◻ under vacuum. Subsequently, 80 μL of methoxyamine (15 mg/mL in pyridine) was added to the residue to react at 30°C for 90 min. Then the residue was derivatized by adding 80 μL N,O-bis-trimethylsilyl-trifluoroacetamide (with 1% trimethylchlorosilane) to react at 70 °C for 60 min.

Finally, derivatized samples were analyzed using the Pegasus HT system (Leco Corporation) coupled with an Agilent 7890B gas chromatography. For each sample, 1 μL volume was injected at a splitless mode with injector temperature at 270 ◻. The flow rate for the carrier gas helium was set at 1.0 mL/min. The oven program started at 70 ◻ for 2 minutes, and then increased to 180 ◻ with a rate of 10 ◻/min, to 230 ◻ with a rate of 6 ◻/min, finally to 295 ◻ with a rate of 40 ◻/min. The temperature of 295 ◻ was held for 5 minutes. The temperatures of transferline interface and ion source were set at 270 ◻ and 220 ◻ respectively. Of note, quality control samples were prepared with a mix of 17 standards and injected at a regular interval to monitor the data quality.

Data were obtained with a m/z range from 50 to 550 at an acquisition rate of 20 spectra per second. Raw data were undergone pre-treatment of baseline correction, de-noising, smoothing, alignment, time-window splitting, and multivariate curve resolution. Metabolites were identified by comparison with the internal library built with the standard reference chemicals. Particularly, a previously reported approach for mass spectral deconvolution [42] was used to determine the peak identities. To validate the identities of differential metabolites, raw chromatograms and true mass spectrometry spectra of representative differential metabolites, as well as the mass spectrometry spectra in our reference library were showed in Supplemental File for Identifying Metabolites.

### Immunohistochemistry staining and western blot assays

Immunohistochemistry (IHC) staining was conducted according to our previous description [41]. Tissue samples were stained with antibody against keratin 14 (cat#ab7800, Abcam, Cambridge, United Kingdom), Ki-67 (cat# ab245113, Abcam, Cambridge, United Kingdom) or nonspecific IgG as negative control.

Western blot was performed as previously described [40, 41, 43]. Cells cultured *in vitro* were digested with 0.25% trypsin and lysed using RIPA buffer (cat#R0278, Sigma-Aldrich, St Louis, MO) containing 1% (vol/vol) protease inhibitor cocktail (cat#P8340, Sigma-Aldrich, St Louis, MO) on ice. Supernatants of cell lysates containing total proteins were obtained by centrifugation at 12,000 ×g for 10 minutes and their concentrations were determined by using a BCA assay kit (cat#23225, Thermo Scientific, Waltham, MA). Protein extracts mixed with sample loading buffer (cat#1610747, Bio-rad, Hercules, CA) were boiled for 10 minutes, resolved by SDS-PAGE, and then transferred to PVDF membranes. After incubation with primary antibodies individually overnight at 4°C, including proliferating cell nuclear antigen (PCNA) (cat#2586S, Cell Signaling Technology, Boston, MA), H2A histone family member X (H2AX)(cat#7631S, Cell Signaling Technology, Boston, MA), phospho-histone H2AX (cat#9718S, Cell Signaling Technology, Boston, MA), E-cadherin (cat#3195T, Cell Signaling Technology, Boston, MA), N-cadherin (cat#13116T, Cell Signaling Technology, Boston, MA), and Vimentin (cat#5741T, Cell Signaling Technology, Boston, MA), the membranes were washed and then incubated with secondary antibody conjugated with IgG-HRP (cat#7074, Cell Signaling Technology, Boston, MA). Images were captured by ChemiDoc Touch Imaging System (Bio-rad, Hercules, CA). The original gels for western blot assays were provided in **Figure S5-S8**.

### Biological function assays of pipecolic acid and 1-oleoylglycerol

Two human ESCC cell lines, KYSE150 (Stem Cell Bank, Chinese Academy of Sciences) and Eca109 (Stem Cell Bank, Chinese Academy of Sciences), were enrolled in this study. Firstly, the biological functions of pipecolic acid were investigated. Cells were seeded and grown to 80% confluence in 96-well plates for ROS measurement and cell viability analysis, or in 6-well plates for analysis of H2AX and γ-H2AX and 8-oxo-dG production. Cells were then pretreated with or without pipecolic acid (cat#HY-W012734, MedChemExpress, Princeton, NJ) in complete medium for 12 hours. Subsequently, Rosup (cat#S0033S, Beyotime, Shanghai, China) was added into the medium at a concentration of 0.2 mg/mL for 6 hours, while H_2_O_2_ (cat#l1522097, Aladdin, Shanghai, China) was added into the medium at a concentration of 1 μM for 4 hours. For analysis of ROS and cell viability, cells were treated in the plates using the corresponding kits. For analysis of H2AX and γ-H2AX and 8-oxo-dG production, cells were digested with trypsin and harvested for measurement.

In addition, for exploration of the impact of pipecolic acid on cell migration, transwell assay was conducted and pipecolic acid was added into the upper and lower chambers at a concentration of 2.5 mM. Furthermore, for investigation of the influence of pipecolic acid on EMT protein marker expression, cells were seeded in 6-well plate, treated with 2.5 mM pipecolic acid for 24 hours, and harvest after trypsin digestion for western blot assay.

Secondly, the biological functions of 1-oleoylglycerol (cat#M7765, Sigma-Aldrich, St Louis, MO) were studied. Cells were seeded in 96-well plates. For cell viability assay, different concentrations of 1-oleoylglycerol as indicated in the figure were added into the wells. After 72 hours, cell viability was measured. For ROS generation assay, 100 μM 1-oleoylglycerol were added into the wells. After treatment for 4 hours, intracellular ROS generation was assessed.

### Measurement of reactive oxygen species, 8-oxo-dG, cell viability, and cell migration

Production of reactive oxygen species (ROS) was measured by 2’,7’-dichlorofluorescein diacetate (DCFH-DA, cat# S0033S-1, Beyotime, Shanghai, China). Briefly, ESCC cells (1×10^4^/well) were incubated in medium containing 10 μM DCFH-DA for 30 minutes at 37◻, then washed with serum-free medium three times. The fluorescence was analyzed using the fluorescence spectrophotometer (Excitation 488 nm, emission 525 nm).

Generation of 8-oxo-dG was assessed by using a modified ELISA kit (Cat#JL11850, Jianglaibio, Shanghai, China) according to the manufacturer’s instructions. Briefly, ESCC cells were crushed by ultrasound, centrifuged at 1,200 × g for 10 minutes, and the supernatant was collected for detection. For each sample, 50 μL supernatant was added to each well precoated with 8-oxo-dG trapping antibody, and then 100 μL horseradish peroxidase-labelled antibody was added into each well. At the meantime, calibration curve was executed by adding distinct concentrations of standard compound solution into wells (50 μL/well) with the same treatment protocol. Next, the 8-oxo-dG measurement plate was incubated at 37◻ for 1 hour. After washing for 5 times, 100 μL tetramethylbenzidine was added to each well and the measurement plate was incubated at 37 ◻ for 15 minutes in the dark. Subsequently, 50 μL reaction termination solution was added to each well and mixed thoroughly. Finally, optical density values at 450 nm of wells of the measurement plate were recorded for quantifying 8-oxo-dG levels.

Cell viability was assessed by using Cell Counting Kit-8 (cat# CK04, Dojindo, Kumamoto, Japan) according to the manufacturer’s instructions.

Cell migration was analyzed in Corning Transwell Permeable Supports with polycarbonate membranes (24-well plate, 8 μm pore size) (cat#18320044, Corning, Corning, NY). ESCC cells were harvested after trypsin digestion, rinsed twice in PBS, resuspended in RPMI 1640 medium with 2.5 mM pipecolic acid or without pipecolic acid as indicated in figure, and then seeded into the upper chambers of the transwell plates at a density of 1 × 10^5^/well. The lower chamber of each well was filled with complete medium with 20% fetal bovine serum containing 2.5 mM pipecolic acid or no pipecolic acid as indicated in figure. After incubation in the cell incubator for 24 hours, the migrating ESCC cells were observed by crystal violet staining.

### Statistical and bioinformatic analyses

To decipher the difference in serum metabolomic profiles between HCs and preoperative ESCC patients, or between preoperative and postoperative ESCC patients, a supervised multivariate regression model orthogonal partial least squares-discriminant analysis (OPLS-DA) was constructed with the software SIMCA-P+ (version 11.0, Umetric, Umea, Sweden). To understand the difference in global metabolism of sera among control mice, 4-NQO-induced mice for 16 weeks and 4-NQO-induced mice for 28 weeks, supervised partial least-squares (PLS) regression was fitted using the software SIMCA-P+. The model parameters R^2^Y and Q^2^ were recorded. R^2^Y indicated the ‘goodness of fit’ in the data. Q^2^, calculated by a cross-validation procedure, indicated the predictability of the model.

Subsequently, univariate statistical analysis was performed to identify significantly altered metabolites. For the metabolomic data of human sera, nonparametric Wilcoxon rank-sum test with Bonferroni adjustment was used to compare each two groups. For the metabolomic data of control mice, 4-NQO-induced mice for 16 weeks and 4-NQO-induced mice for 28 weeks, one-way analysis of variance (ANOVA) was executed and post hoc test for multiple comparisons was implemented using least significant difference (LSD) method. For the data from cell experiments, Student’s t test was used for statistical analysis. All univariate tests were 2-sided, and *P* values < 0.05 were considered statistically significant. Differential metabolites were displayed by heat maps, which were generated by using a R package ComplexHeatmap [44].

Metabolite set enrichment analysis (MSEA) for the differentially expressed metabolites was performed using a web-based tool MetaboAnalyst 4.0 [45, 46] and algorithm of quantitative enrichment analysis was selected. Pathway-associated metabolite sets (SMPDB) was selected as the metabolite set library for MSEA analysis.

## Ethical statement

All participants provided informed written consent in accordance with the regulation of the Institutional Review Boards of the Affiliated Tumor Hospital of Nantong University (Approval number 2019-022) in agreement with the Declaration of Helsinki. Mouse study was approved and conducted by Wuhan Servicebio Technology Co., Ltd (Approval number 2018015).

## Data availability

The mass spectrometry data of serum metabolome of healthy volunteers, ESCC patients and mice have been deposited in The National Omics Data Encyclopedia (NODE) database at Bio-Med Big Data Center affiliated with Shanghai Institute of Nutrition and Health, Chinese Academy of Sciences (Project ID: OEP001955), and are publicly accessible at https://www.biosino.org/node/project/detail/OEP001955.

## CRediT author statement

**Wen-Lian Chen**: Conceptualization, Supervision, Formal analysis, Data Curation, Visualization, Writing–Original Draft, Writing–Review & Editing. **Xiaoxia Jin**: Conceptualization, Formal analysis. **Xin Jin**: Conceptualization, Supervision, Validation. **Lei Liu**: Resources, Investigation, Supervision. **Jia Wu**: Validation, Formal analysis, Data Curation, Writing–Original Draft, Visualization. **Minxin Shi**: Resources. **Fengying Wang**: Resources. **Haimin Lu**: Resources. **Jibing Liu**: Resources. **Weiqin Chen**: Resources. **Guanzhen Yu**: Investigation, Formal analysis. **Dan Liu**: Investigation. **Jing Yang**: Investigation. **Qin Luo**: Investigation. **Yan Ni**: Investigation, Data Curation.

## Competing interests

The authors declare that they have no financial competing interests.

### Acknowledgments

This work was supported by National Natural Science Foundation of China (31970708, 81770147, 81802891), The National Scientific and Technological Major Special Project of China (2019ZX09201004-002-013), National Thirteenth Five-Year Science and Technology Major Special Project for New Drug Innovation and Development (2017ZX09304001), Research fund of Shanghai Municipal Commission of Health (20174Y0090), Shanghai Rising-Star Program (18QA1404100), Program for Professor of Special Appointment (Eastern Scholar) at Shanghai Institutions of Higher Learning, Shanghai Youth Talent Program, Shanghai Municipal Key Clinical Specialty (shslczdzk03701), The Three-Year Plan of Shanghai Municipality for Further Accelerating The Development of Traditional Chinese Medicine (ZY(2018-2020)-CCCX-1016), Shanghai Chenguang Program (18CG47), The grant from Nantong Tumor Hospital (BS201909), Gaofeng Clinical Medicine Grant of Shanghai Municipal Education Commission, Health Commission of Pudong New Area Health and Family Planning Scientific Research Project (PW2019E-1), and Xinling Scholar Program of Shanghai University of Traditional Chinese Medicine.

