## Supplemental information for "New metabolic alterations and predictive marker pipecolic acid in sera for esophageal squamous cell carcinoma"

**Table S1.** The relative standard deviation (RSD) values of standard chemicals in quality control (QC) samples and the mean RSD for all the tested standard chemicals in the study.

| Chemicals | RSD (%) | Quality control check |
| --- | --- | --- |
| 11,14-Eicosadienoic acid | 3.83 | passed |
| 4-Hydroxybutyric acid | 2.57 | passed |
| Alpha-Tocopherol | 6.13 | passed |
| Aspartic acid | 2.34 | passed |
| Cholesterol | 1.97 | passed |
| Fructose | 1.17 | passed |
| Glutamic acid | 2.66 | passed |
| Glycine | 5.28 | passed |
| Leucine | 1.26 | passed |
| Nicotinic acid | 1.07 | passed |
| Palmitoleic acid | 2.15 | passed |
| Proline | 2.09 | passed |
| Ribose | 1.56 | passed |
| Stearic acid | 1.51 | passed |
| Thymine | 3.23 | passed |
| Urea | 1.40 | passed |
| Valine | 2.12 | passed |
| Mean RSD (%) | 2.49 |  |

**Table S2. Composition of the 10% amino acid solution.**

| Ingredient | Density in solution, g/1000mL |
| --- | --- |
| Isoleucine | 8.80 |
| Lysine acetate | 10.60 |
| Phenylalanine | 1.60 |
| Tryptophan | 1.50 |
| Arginine | 8.80 |
| Glycine | 6.30 |
| Proline | 7.10 |
| Asparagine | 0.55 |
| N-acetyl-L-cysteine | 0.80 |
| N-acetyl-L-tyrosine | 0.86 |
| Leucine | 13.60 |
| Methionine | 1.20 |
| Threonine | 4.60 |
| Valine | 10.60 |
| Histidine | 4.70 |
| Alanine | 8.30 |
| Aspartic acid | 2.50 |
| Ornithine hydrochloride | 1.66 |
| Serine | 3.70 |
| Glutamate | 5.70 |

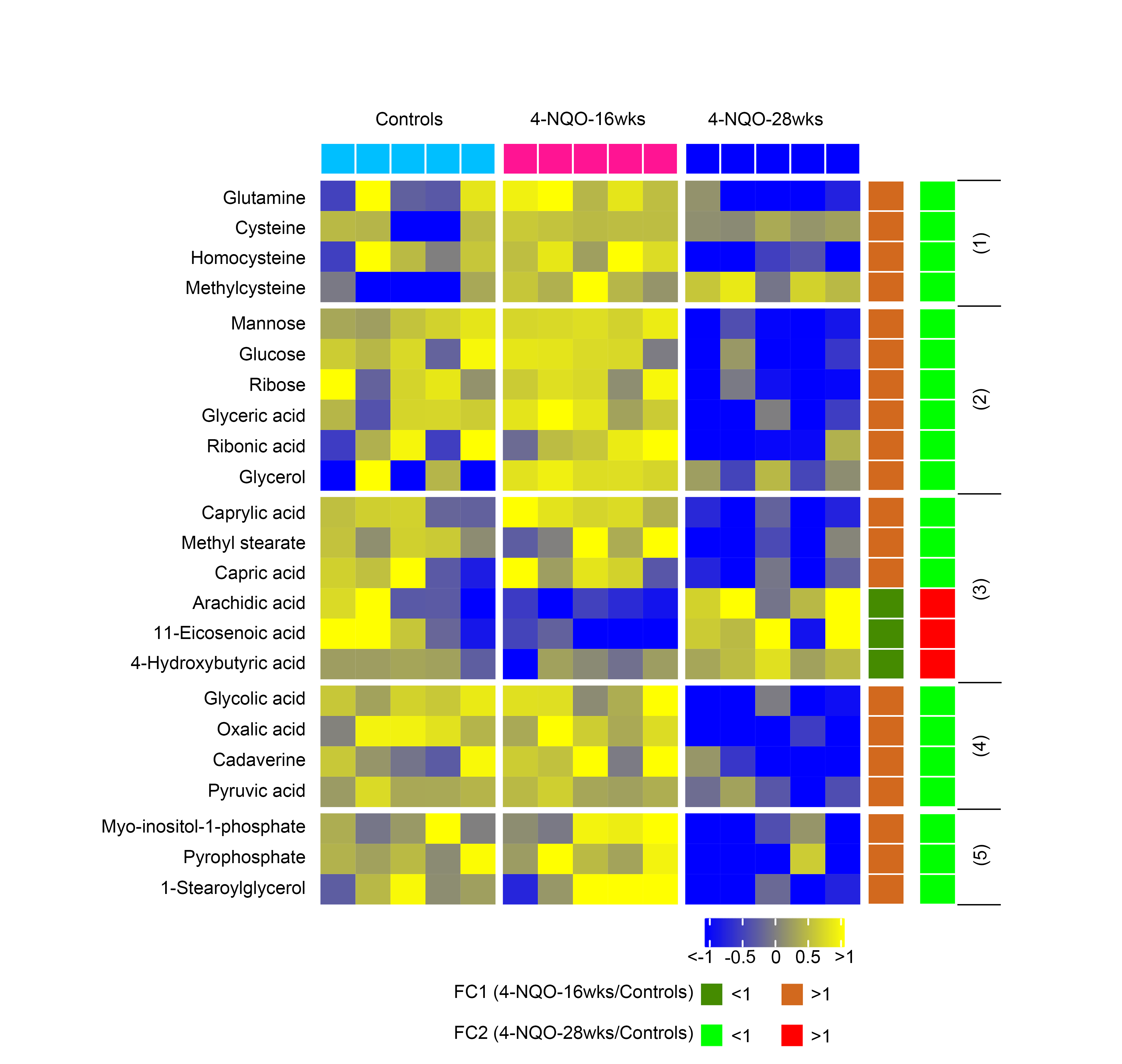

**Figure S1. Heat map displaying differential serum metabolites among control, dysplasia and cancerization mice with no progressive changes.**

The metabolites are subclassified as follows: (1) Amino acids, (2) Carbohydrates, (3) Lipids including fatty acids, (4) Organic acids, (5) Unclassified.

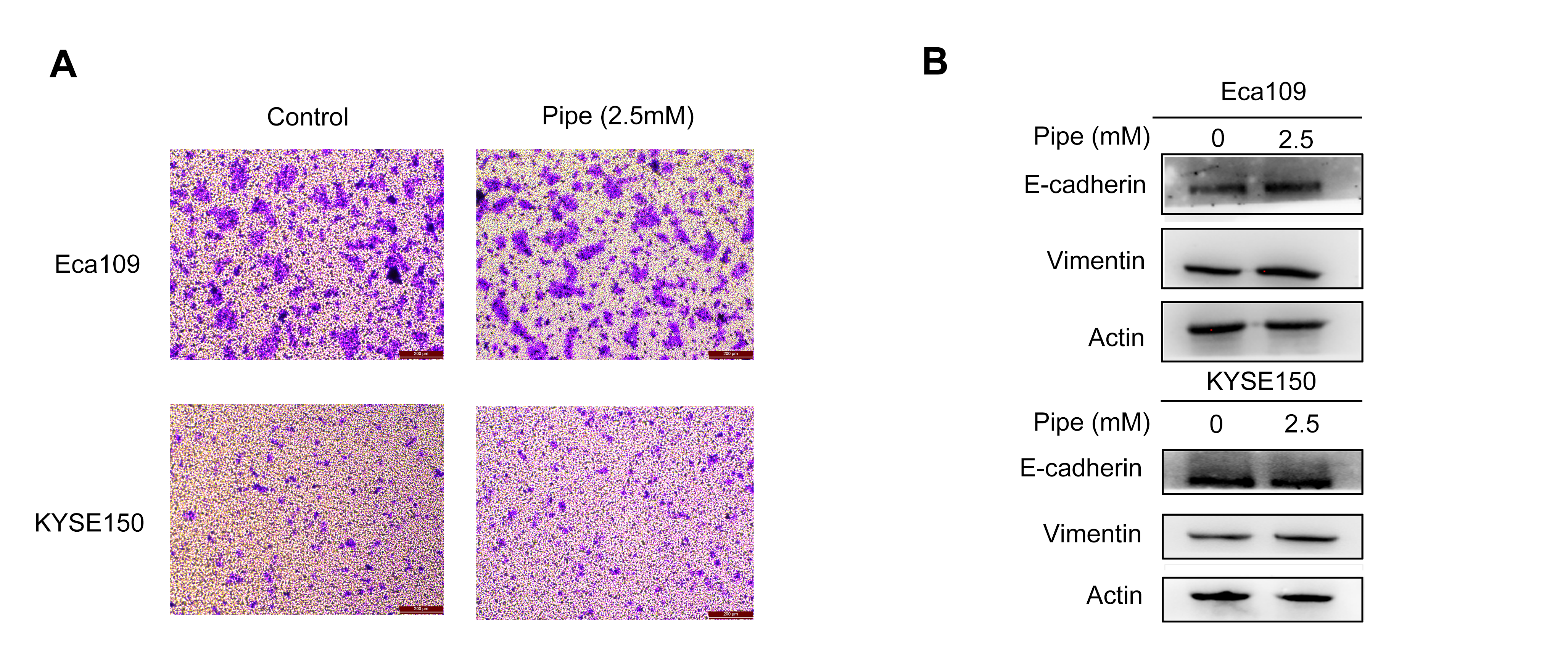

**Figure S2. The impact of pipecolic acid on ESCC cell migration and expression of EMT protein markers.**

**A,** Transwell assay revealing the influence of pipecolic acid on ESCC cell migration.

**B,** Western blot showing the impact of pipecolic acid on the expression of EMT protein markers in ESCC cells.

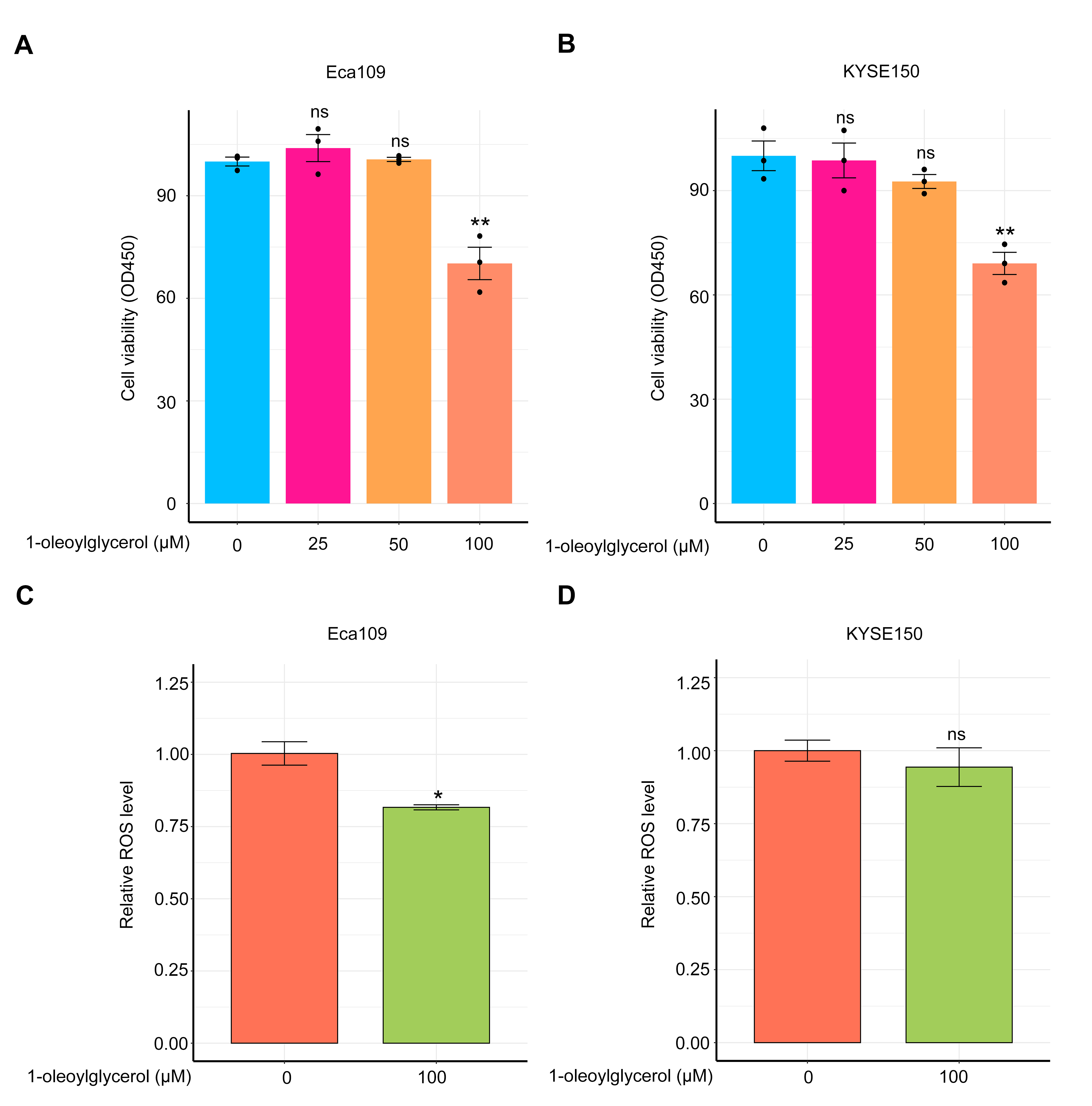

**Figure S3. The impact of** **1-oleoylglycerol on ESCC cell growth and ROS generation.**

**A,** The inhibitory effect of 1-oleoylglycerol on ESCC cell growth.

**B,** The influence of 1-oleoylglycerol on ROS production of ESCC cells.

Error bars represent mean ± SEM. ^*^*P* < 0.05, ^**^*P* < 0.01, ^***^*P* < 0.001, by comparison with the group treated with 0 μM 1-oleoylglycerol, 2-tailed Student’s t test.

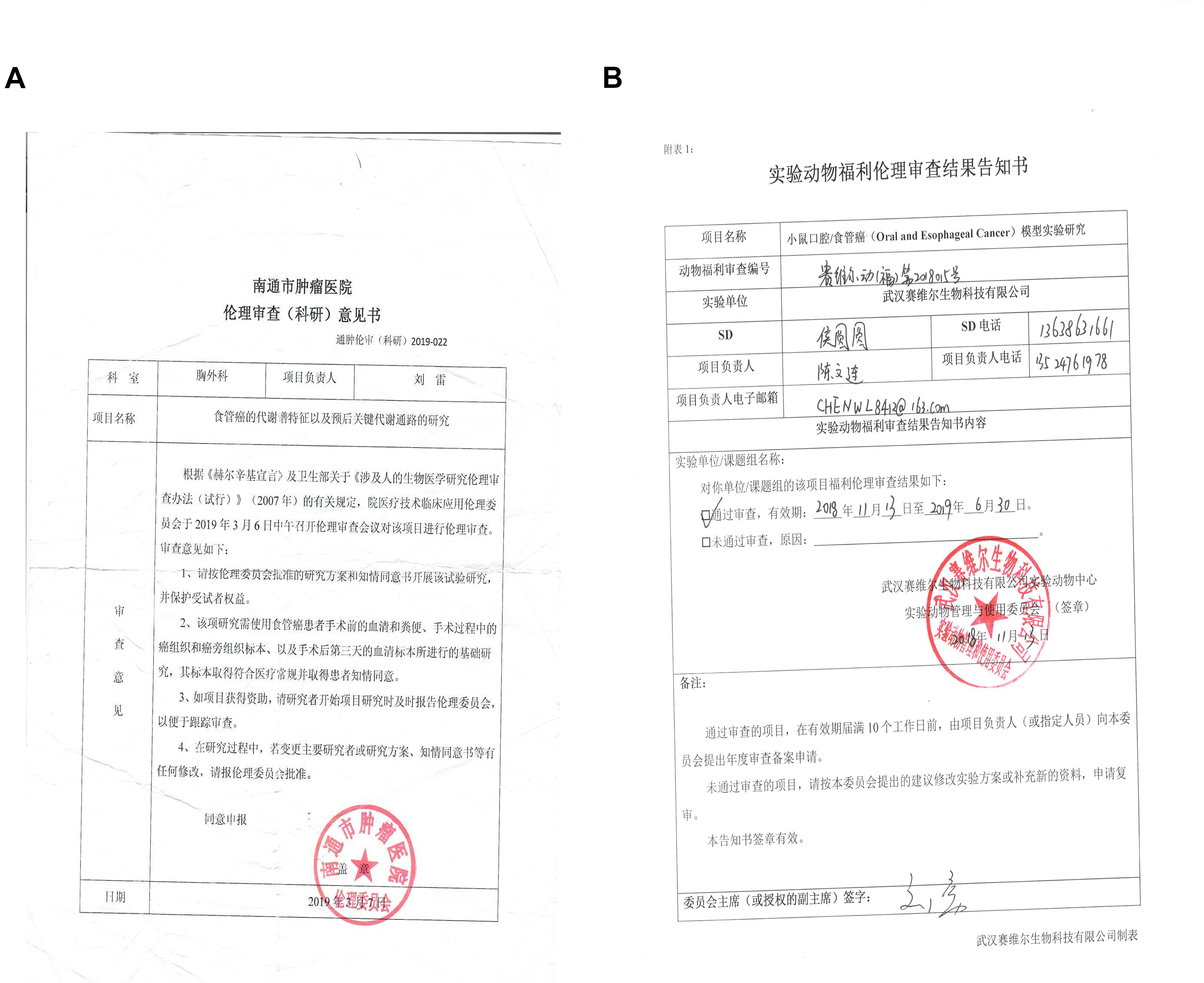

**Figure S4.** **The photocopies of the ethic approval for ESCC patient enrollment and mouse experiment.**

**A,** The photocopy of the ethic approval for ESCC patient enrollment.

**B,** The photocopy of the ethic approval for mouse experiment.

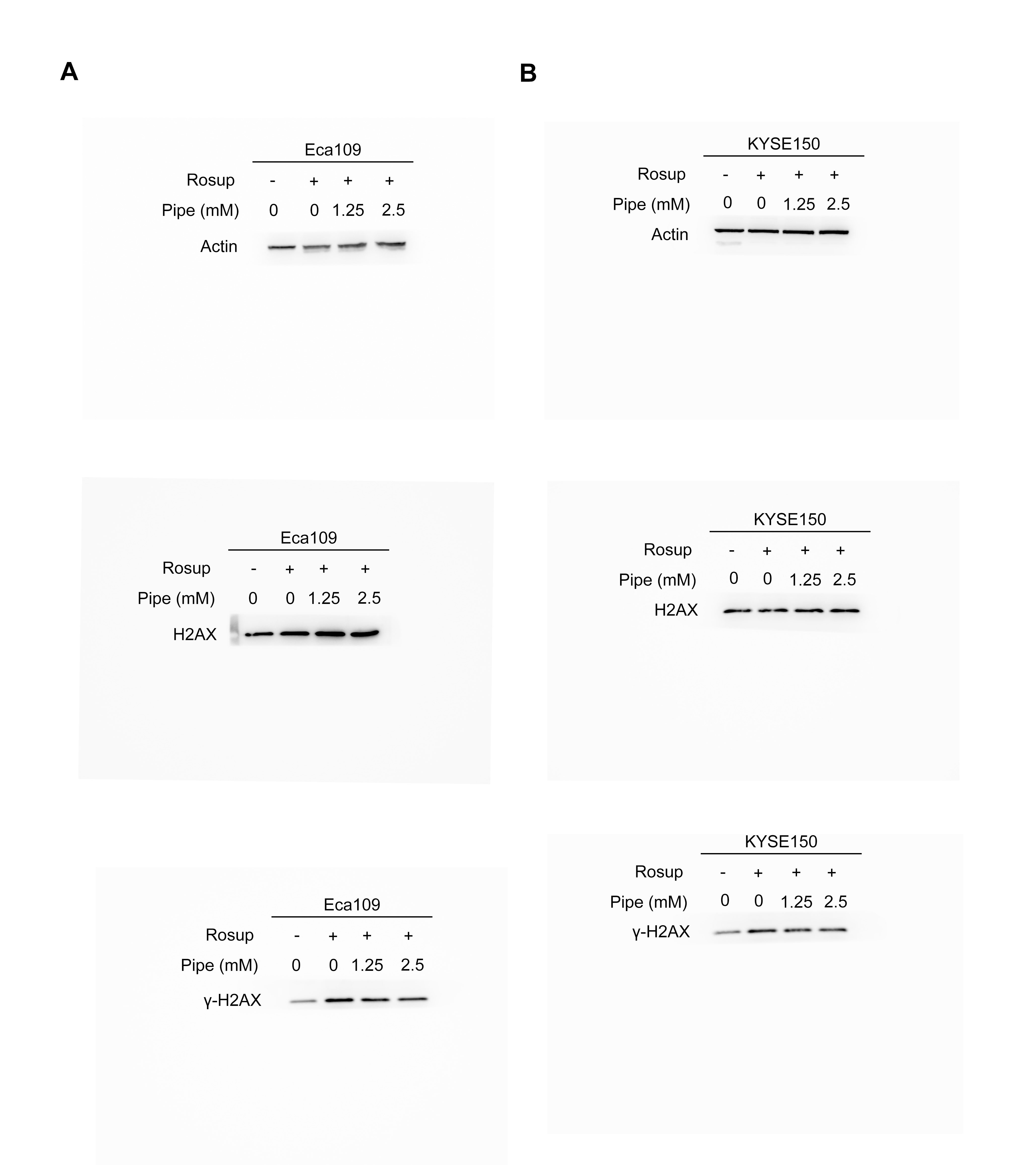

**Figure S5. The original gels for western blot assays in Figure 7C.**

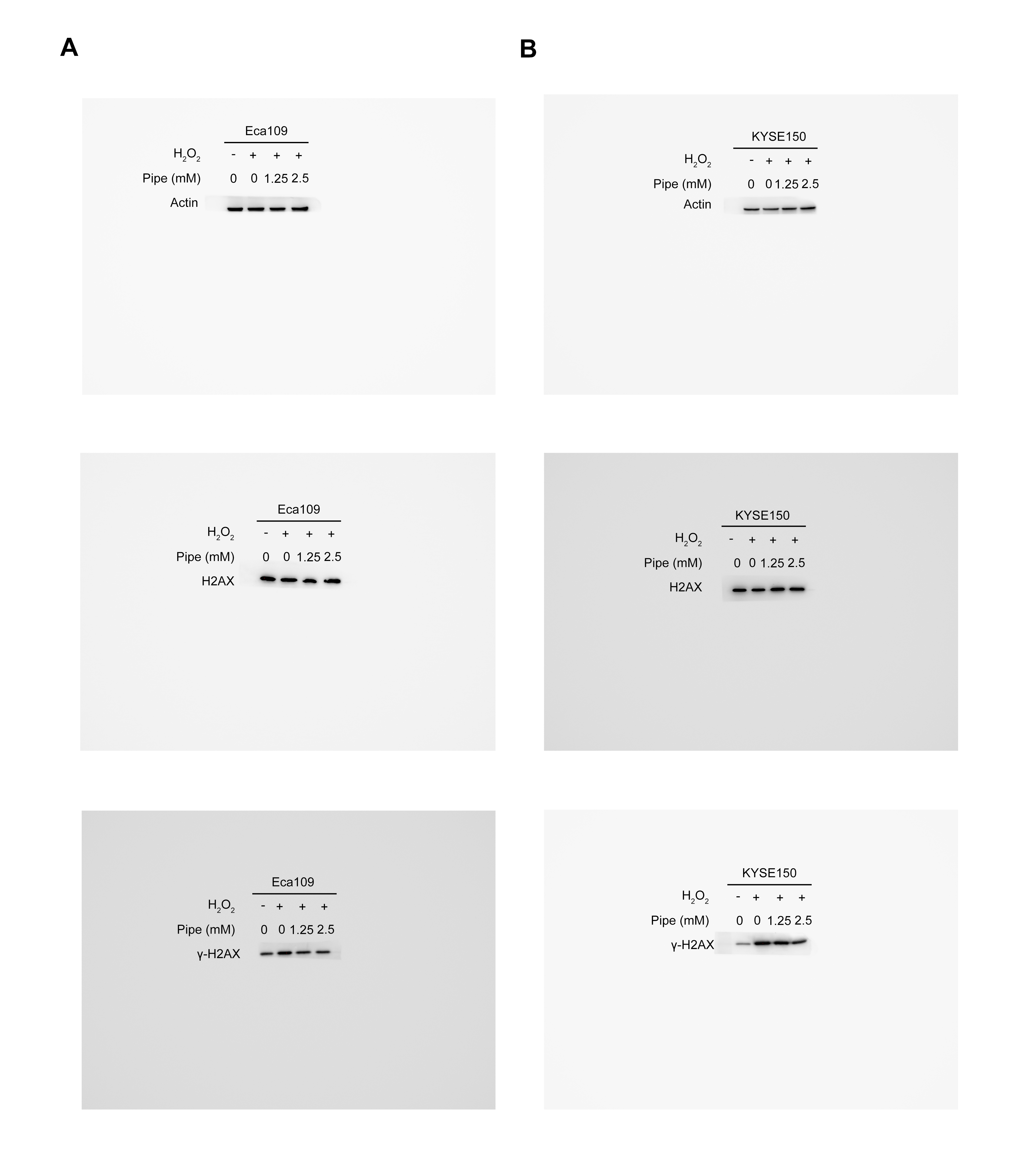

**Figure S6. The original gels for western blot assays in Figure 7D.**

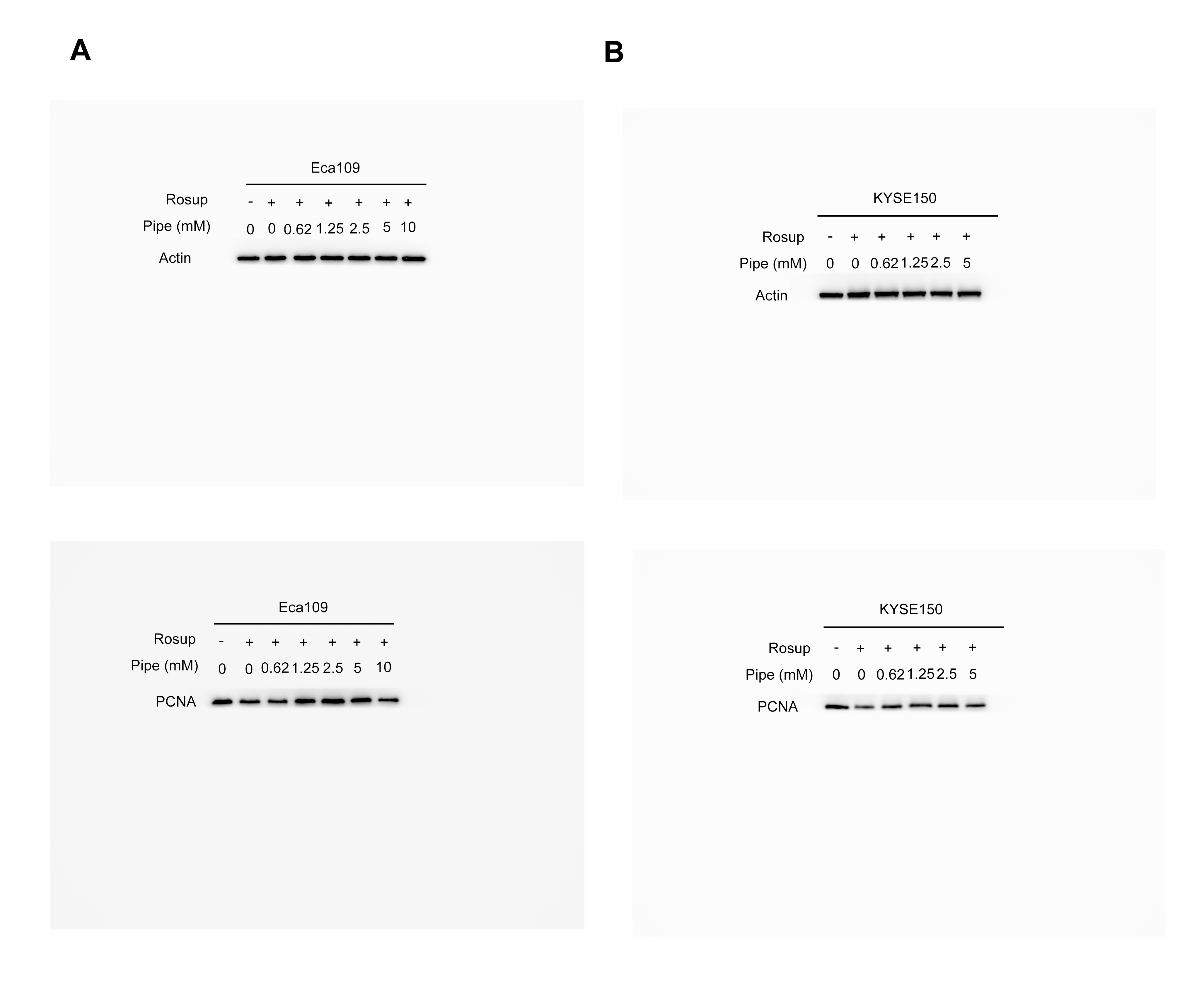

**Figure S7. The original gels for western blot assays in Figure 7G.**

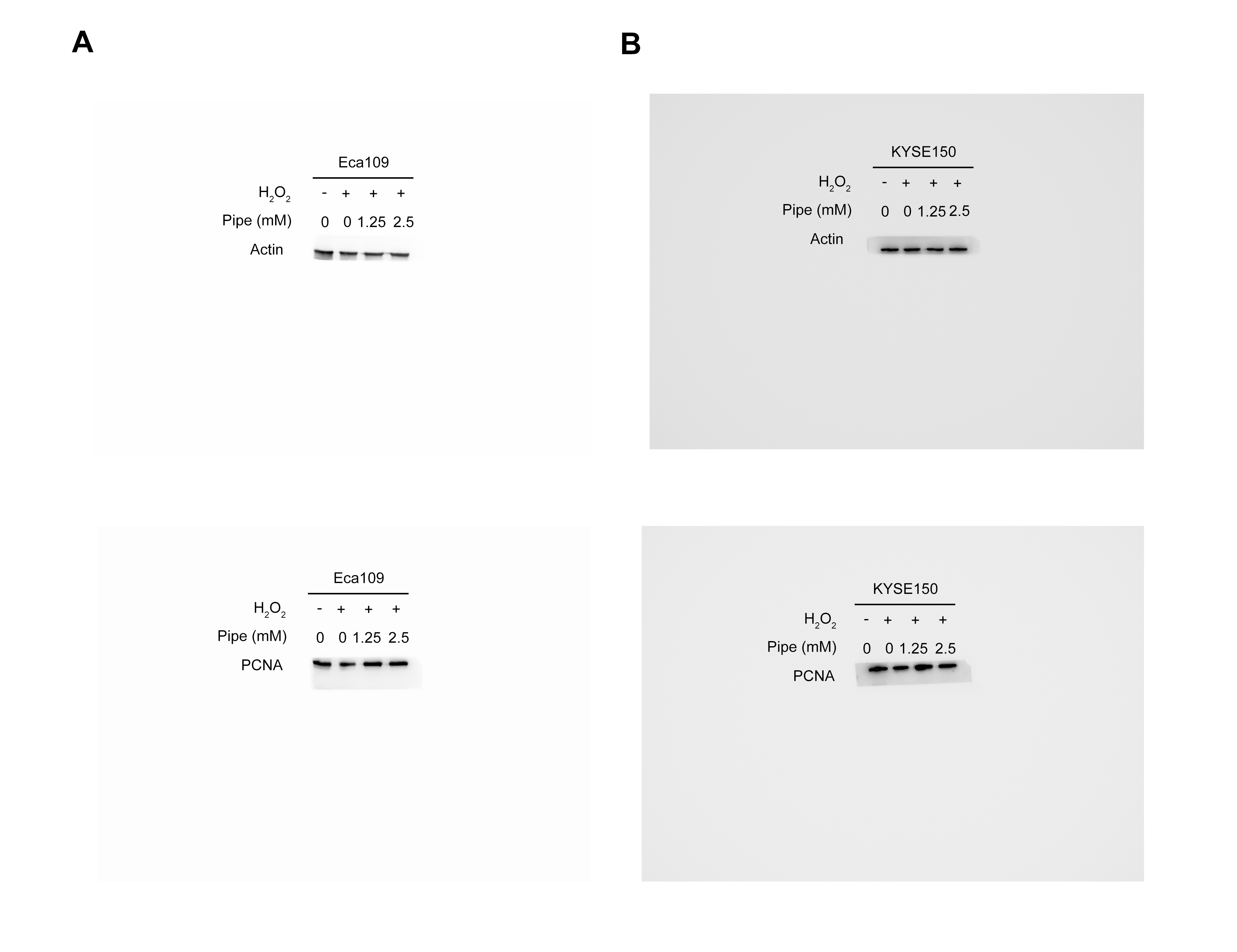

**Figure S8. The original gels for western blot assays in Figure 7H.**

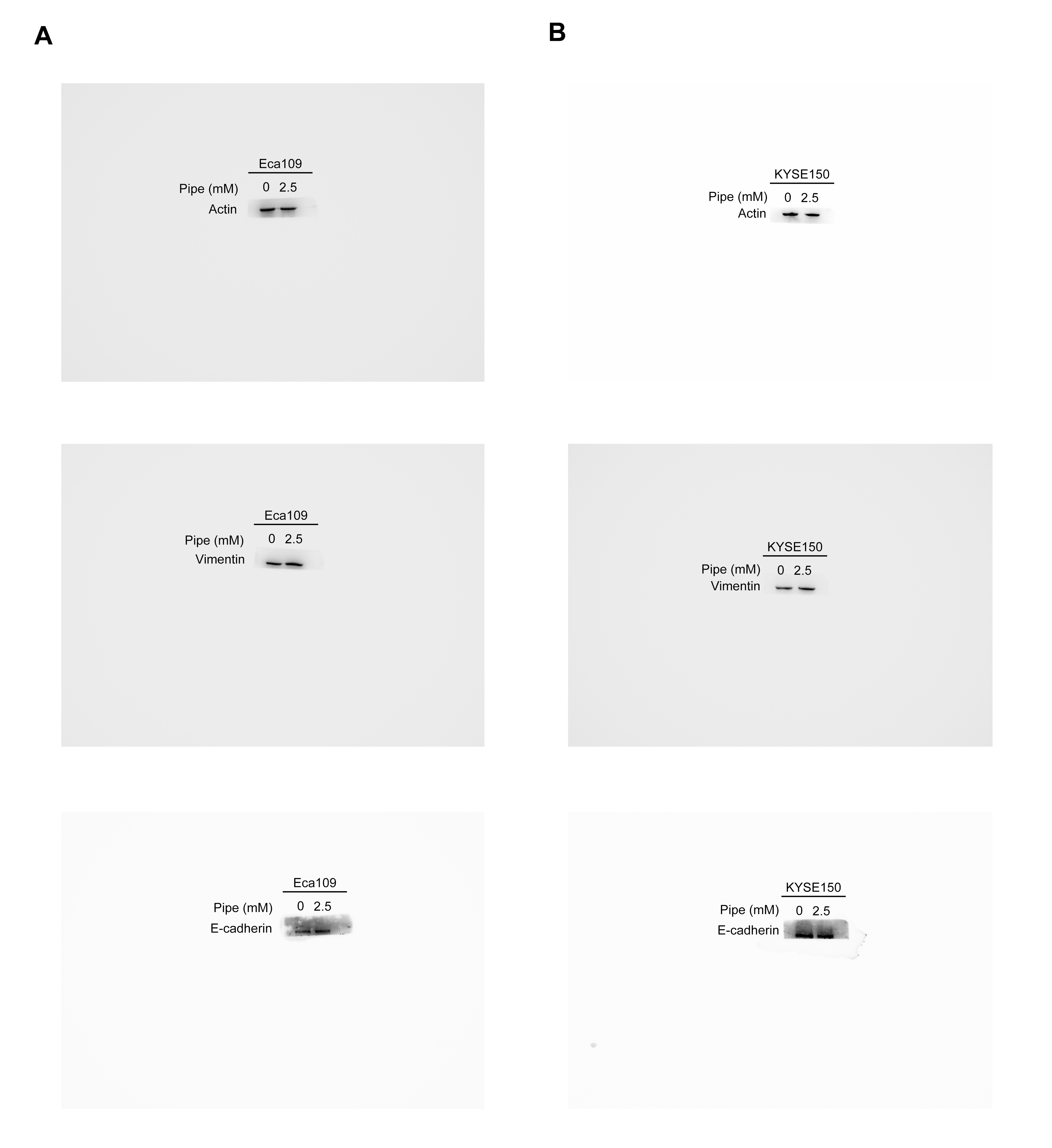

**Figure S9. The original gels for western blot assays in Figure S2.**
